# Optimized LC-MS method for simultaneous polyamine profiling and ADC/ODC activity quantification and evidence that ADCs are indispensable for flower development in tomato

**DOI:** 10.1101/2025.07.09.663906

**Authors:** Erin Samantha Ritchie, Edda von-Roepenack-Lahaye, Dennis Perrett, Dousheng Wu, Thomas Lahaye

**Affiliations:** University of Tübingen, ZMBP - General Genetics, Auf der Morgenstelle 32, 72076 Tuebingen, Germany; University of Tübingen, ZMBP – Central Facilities, Auf der Morgenstelle 32, 72076 Tuebingen, Germany; Hunan Key Laboratory of Plant Functional Genomics and Developmental Regulation, College of Biology, Hunan University, Changsha 410082, China

## Abstract

Polyamines (PAs) are essential for plant development and stress responses, requiring tight homeostatic regulation. Many PA enzymes are regulated post-transcriptionally, making traditional transcript-based methods ineffective in determining their abundance, highlighting the need for alternative approaches to study PA homeostasis. Here, we refined a liquid chromatography-mass spectrometry (LC-MS) based method to simultaneously quantify activities of two key PA synthesizing enzymes – arginine decarboxylase (ADC) and ornithine decarboxylase (ODC) – from plant tissues using stable isotope substrates. By optimizing substrate concentrations, we increased assay sensitivity >10-fold in tomato leaf tissue. We further adapted this protocol for *Nicotiana benthamiana*, a model plant widely used for transient recombinant protein expression. Expression of epitope-tagged ADCs in this system revealed a direct correlation between protein abundance and enzymatic activity, demonstrating that ADC activity can infer its protein abundance in native tissues. Proof-of-principle experiments with the *N. benthamiana* expression system, confirm substrate specificity of tomato ADC and ODC enzymes and essential catalytic residues of tomato ADCs. Beyond enzymatic activities, our LCMS-based method also permits quantification of 11 PA network metabolite concentrations from the same LCMS sample. Visualizing this data as a heatmap pathway diagram, alongside ADC/ODC activities provides a comprehensive overview of PA metabolism in plant tissues. We also studied tomato CRISPR-Cas9-induced mutants deficient in ADC or ODC, complemented by phenotypic analysis. LC-MS analysis of an *adc1/adc2* double mutant – an embryo lethal genotype in Arabidopsis – had no detectable agmatine, the product of ADCs. Additionally, despite a reduction in putrescine, no impact on the downstream PAs, spermidine and spermine, was found. The *adc1/adc2* double mutant showed severe developmental abnormalities, including complete flower loss, demonstrating the indispensable role of ADCs in flower development. In summary, our optimized LC-MS approach for simultaneous quantification of ADC/ODC enzyme activity and PA-pathway metabolites, the ability to transiently express and functionally analyze recombinant ADC/ODC proteins *in planta*, and a collection of tomato CRISPR mutants deficient in these enzymes collectively establish a versatile new experimental toolkit to dissect PA homeostasis and PA-dependent developmental processes in plants.

## INTRODUCTION

Polyamines (PAs) are essential metabolites for almost all organisms (Michael, 2016). In plants they are required for various processes such as growth, development, reproduction, senescence, and play key roles in abiotic and biotic stress responses (Alcazar et al., 2010, Chen et al., 2019, Gonzalez et al., 2021, Napieraj et al., 2023, Blázquez, 2024). PAs are low molecular weight compounds consisting of at least two amino groups and exist in various forms: free, bound to cellular moieties (e.g., RNA, DNA, and cell wall components), or conjugated to other metabolites such as hydroxycinnamic acids (Bassard et al., 2010, Chen et al., 2019, Pal et al., 2021). Despite their indispensability and numerous roles, the mechanism by which PA homeostasis is maintained remain unclear.

In plants, PAs are synthesized via two main routes; both converging on the production of putrescine (Figure 1; Liu et al., 2015, Michael, 2016). The first anabolic route involves decarboxylation of arginine into agmatine by arginine decarboxylase (ADC), while the second involves direct decarboxylation of ornithine into putrescine by ornithine decarboxylase (ODC). Putrescine is then converted to the higher PAs, spermidine and spermine. Cadaverine, another PA, is independently synthesized from lysine by lysine decarboxylase (LDC) (Bunsupa et al., 2012, Jancewicz et al., 2016). ADC, ODC, and LDC enzymes belong to the same IV pyridoxyl 5-phosphate (P5P)-dependent decarboxylase family (Sandmeier et al., 1994). These enzymes dimerize head-to-tail configuration, with essential lysine and cysteine residues from each monomer forming two catalytic sites at the interface; the lysine interacting with P5P, and the cysteine responsible for substrate specificity (Poulin et al., 1992, Hanfrey et al., 2001, Bunsupa et al., 2012, Liang et al., 2019).

**Figure 1:**
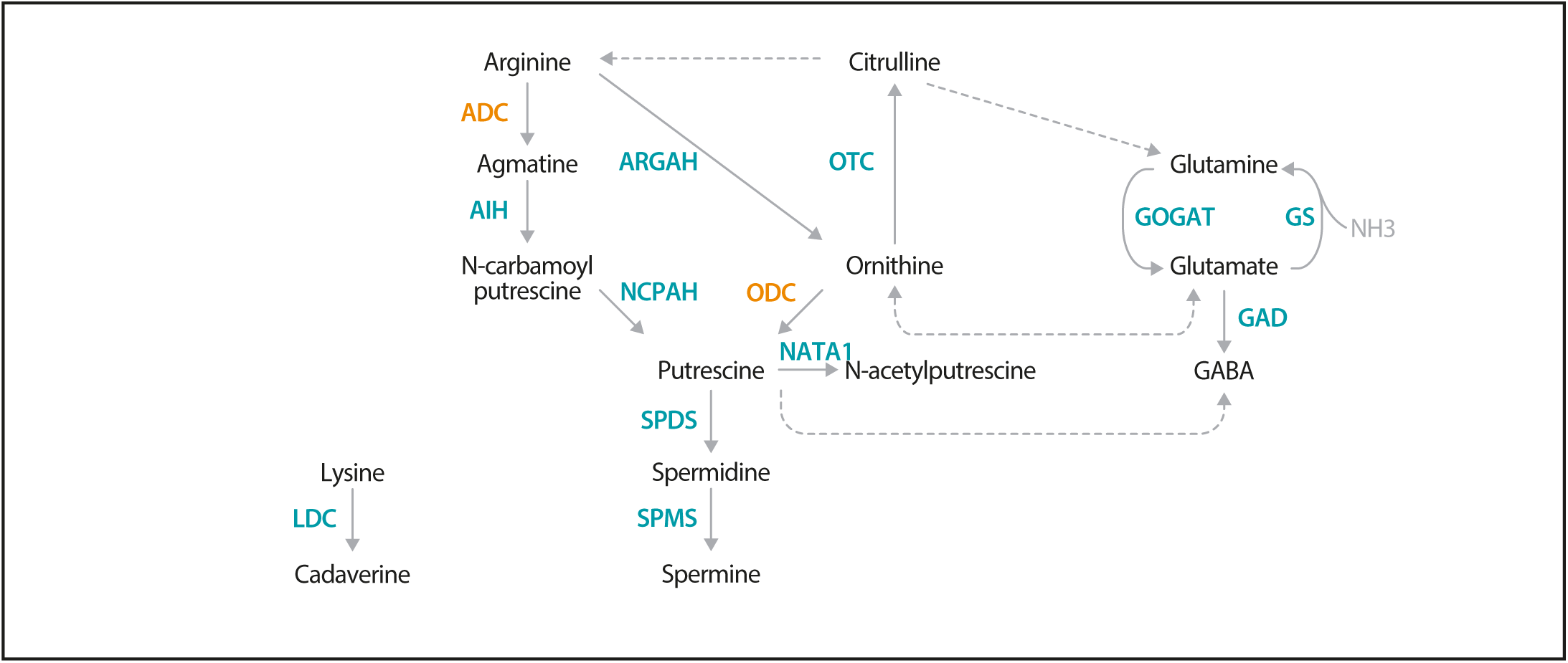
Polyamine metabolic pathway, highlighting the ADC- and ODC dependent pathways. Synthesis of the main plant polyamines (PAs), putrescine, spermidine, and spermine originates with either arginine via the arginine decarbox-ylase (ADC)-dependent route or the ornithine route via ornithine decarboxylase (ODC). Cadaverine, another PA, is produced from lysine via lysine decarboxylase (LDC). Solid arrows represent reactions catalyzed by a single enzyme, while dotted arrows indicate multiple enzymatic steps. Additional enzyme names are as follows: agmatine iminohydrolase (AIH); N-carbamoylputrescine amidohydrolase (NCPAH); spermidine synthase (SPDS); spermine synthase (SPMS); arginase (ARGAH); ornithine-carbamoyl transferase (OTC); glutamate synthase (GOGAT); glutamine synthase (GS); glutamate decarboxylase (GAD); and N-acetyltransferase activity 1 (NATA1).

Although *Arabidopsis thaliana* is a commonly used plant model for genetic and biochemical studies, it is not suitable for investigating canonical PA metabolism which requires both ADC and ODC. This is because many *Brassicaceae* species, including *A. thaliana* lack the PA anabolic enzyme ODC, and therefore rely solely on the ADC-dependent pathway for PA synthesis (Hanfrey et al., 2001, Jiménez-Bremont et al., 2014). As a result, an *A. thaliana adc1/adc2* double mutant is embryonically lethal (Urano et al., 2005), thereby preventing studies in this model plant that could provide insights into which biochemical or developmental processes depend on the functionality of ADC1/2 proteins.

In contrast to *A. thaliana* and some other Brassicaceae species, solanaceous species (e.g., tomato [*Solanum lycopersicum*; *Sl*] and *Nicotiana benthamiana*) possess the canonical PA biosynthetic pathway found in most land plants, making them ideal platforms for studying the mechanistic principles of PA metabolism. In particular, the model plant, *N. benthamiana* provides a unique high-throughput expression platform that enables rapid functional studies of proteins and their mutant derivatives *in planta*, as it can be efficiently transiently transformed using *Agrobacterium tumefaciens* (Ranawaka et al., 2023, Golubova et al., 2024. Moreover, *N. benthamiana* is a host to a wide range of model plant pathogens, as for example the bacterial plant parasites *Pseudomonas* or *Xanthomonas* and therefore provides the unique opportunity to study microbial plant pathogens in a context where PA levels are manipulated by simultaneous *Agrobacterium*-mediated expression of enzymes that induce changes in PA-levels.

For decades, plant PAs have been measured using chromatography-based methods (Richards and Coleman, 1952, Smith and Richards, 1962, Smith, 1970), with recent analytical advancements enabling more accurate PA separation using high-performance liquid or gas chromatography. Subsequent quantification of (often derivatized) metabolites, has been performed using fluorescence or UV detectors, ELISAs, and mass spectrometry (MS; Flores and Galston, 1982, Bouchereau et al., 2000). More recently, methods based on liquid chromatography-mass spectrometry (LC-MS) have been developed for quantification of plant PAs without derivatization, significantly reducing time and cost, and thereby greatly improving the suitability of these approaches for high-throughput analysis (Hakkinen et al., 2007, Sanchez-Lopez et al., 2009). However, most studies solely quantified putrescine, spermidine, and spermine, neglecting pathway precursors and breakdown products. The lack of comprehensive quantitative information on the concentrations of key metabolites involved in anabolic and catabolic PA metabolism represents a major limitation that has hindered a broader understanding of the regulatory principles underlying PA homeostasis.

In addition to studies quantifying PA metabolites, there have also been investigations into the enzymatic activities of enzymes involved in PA homeostasis, with particular focus on the key enzymes ADC and ODC (Smith and Richards, 1962, Smith, 1970, Kaur-Sawhney et al., 1982, Rossi et al., 2018). In the past, the enzymatic activity of ADC and ODC was often determined by supplying radioactive substrates and measuring the release of radioactive CO_2_, rather than quantifying the enzyme-specific metabolic products. However, approaches quantifying ADC/ODC activity based on CO_2_ release are limited, as arginine and ornithine pools are metabolically interconnected (Winter et al., 2015, Majumdar et al., 2016, Joshi and Fernie, 2017). As a result, one labeled substrate can be rapidly converted into another, meaning detection of released radioactive CO_2_ might not be due to the activity of a single enzyme but rather the combined activities of ADC, ODC, and potentially other enzymes involved in PA homeostasis. These considerations highlight that elucidating the regulatory networks underlying PA homeostasis requires analytical methods capable of simultaneously quantifying multiple enzyme activities and PA-related metabolites, methods which are not yet fully established.

Recently, we developed an LC-MS approach to simultaneously quantify ADC activity and the levels of five PA-network metabolites from the same plant sample (Wu et al., 2019, Gallas et al., 2024). To monitor ADC activity specifically, a stable (non-radioactive) heavy isotope variant of arginine (^13^C_6_ arginine) was added to plant extracts, and the formation of the derived heavy isotopic product, ^13^C_5_ agmatine was quantified as a readout of ADC enzymatic activity (Figure 2A). Here, we present our optimized LC-MS-based method, in which the optimization of heavy isotope substrate concentrations, extraction protocols, LC-MS gradient, and metabolite analysis has led to improved sensitivity, a simplified and less laborious workflow, and the ability to detect 11 PAs in a single MS run. We further optimized our assay to enable the simultaneous quantification of both ADC and ODC enzyme activities by using different isotope-labeled metabolites.

**Figure 2:**
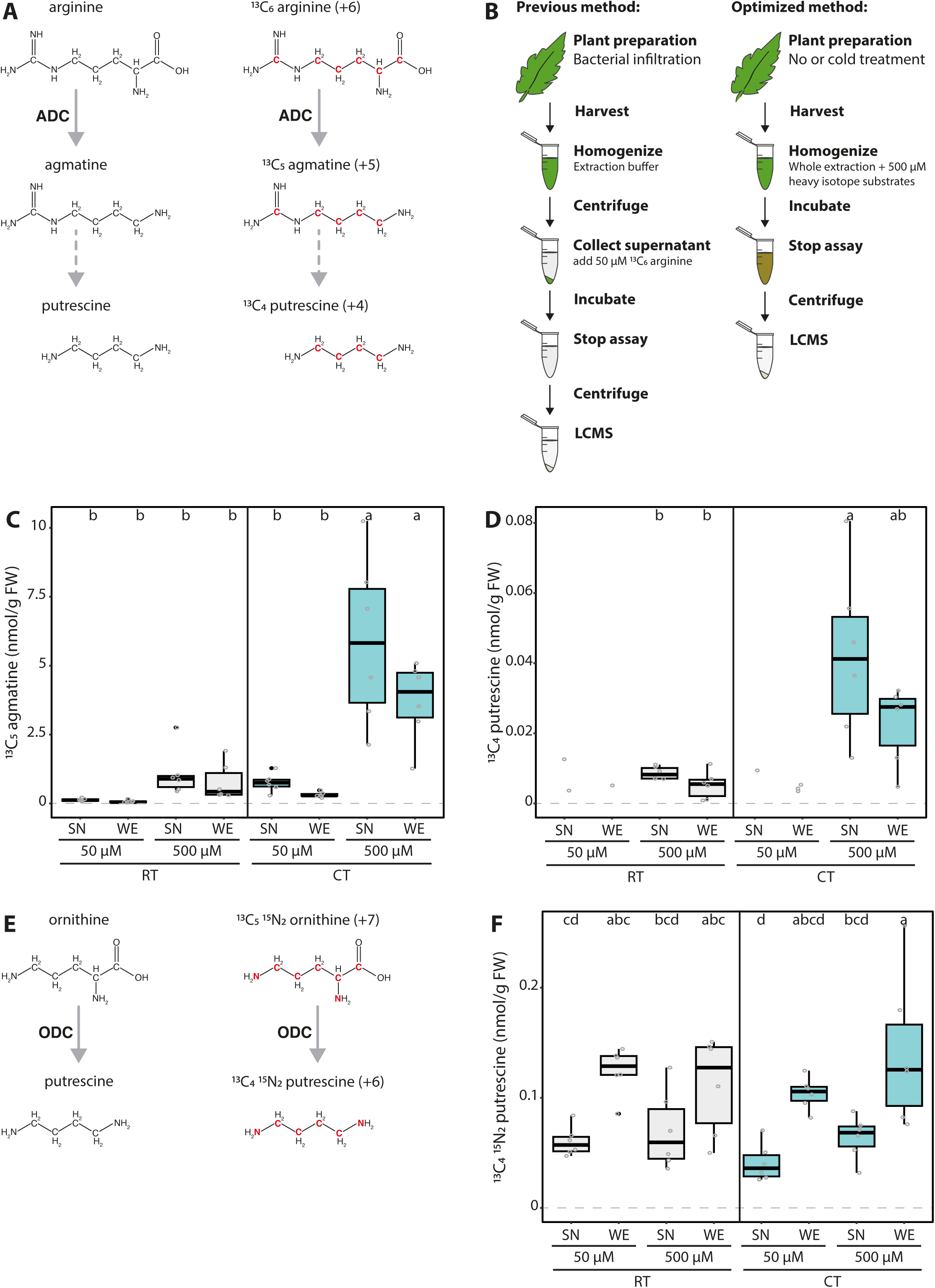
Optimization of LC MS-based method for measuring ADC activity from plant extracts. **A)** Schematic representation of the ADC-catalyzed conversion of natural (left) and heavy (right) isotope arginine variants into agmatine, and the subsequently generated putrescine variant. Red font indicates heavy atoms in these isotopes. **B)** Illustration displaying previously estab-lished LC MS-based quantification of ADC activity according to Wu et al., 2019 (left) and the optimized approach described here (right). In the previously established method, samples of tomato leaves were harvested, flash frozen, ground, homogenized in an extraction buffer, then centrifuged and the supernatant (SN) collected. 50 µM 13C6 arginine is added to each SN sample and incubated for 1 hour at 37 °C before the assay is stopped by adding 5 µL of heptafluorobutyric acid (HFBA) and centrifuged (30 minutes, 4 °C). In the optimized protocol, no initial centrifugation step is required; instead, heavy isotope substrates (500 µM) are added directly to the buffer used to homogenize the whole extracts (WE). **C)** ADC activity is increased upon higher substrate concentration but not impacted by extraction procedure. ADC activity (13C5 agmatine in nmol/g of fresh weight [FW] tissue) in tomato leaf samples, pretreated either with room temperature (RT; grey) or cold treatment (CT; blue), with either 50 or 500 µM 13C6 arginine added to the SN or WE. **D)** The synthesis of heavy putrescine is only detectable at high substrate concentrations. 13C4 putrescine (nmol/g FW), generated from ADC-produced 13C5 agmatine in the same samples presented in C). **E)** Graphical representation of the ODC-catalyzed conversion of natural (left) and heavy (right) isotope ornithine variants into putrescine. Red font indicates heavy atoms in these isotopes. **F)** ODC activity can successfully be simultaneously measured using LC-MS method. ODC activity (13C415N2 putrescine in nmol/g FW) in the same samples presented in C) and D). For C, D, and F, all concentrations were normalized to the internal D5 tryptophan control. Statistical significance was determined using ANOVA followed by Tukey’s post hoc HSD test; different letters above each box indicate statistically significantly groups (a = 0.05). Each biological replicate is represented by a grey circle; n = 6. If a metabo-lite could not be detected in all samples, only the remaining replicates are displayed without a boxplot and this group was removed from the statistical analysis.

This optimized protocol, originally established using tomato leaf extracts, was not suitable for analyzing extracts from the model plant *N. benthamiana*, yet by incorporating an additional extraction step, it is now compatible with *N. benthamiana* leaf extracts. Using the *N. benthamiana* transient expression system in conjunction with these optimized assay conditions, we verified that ADC activity directly correlates to ADC protein abundance and corroborated essential tomato ADC (SlADC) residues for catalytic activity as well as ADC/ODC substrate specificity. Finally, we showcase our LC-MS method by conducting ADC activity and PA profiling on four CRISPR-Cas9-generated tomato mutants, including an *adc1/adc2* double mutant, and characterize their phenotypes. Through this, we demonstrate the importance of ADCs for PA metabolism and their crucial role in plant and flower development.

## MATERIALS AND METHODS

### Plant materials and growth conditions

Tomato (Moneymaker) and *N. benthamiana* were grown at 21 ± 3 °C, 30-50% humidity, and a 16/8-hour light/dark photoperiod.

### CRISPR**-**Cas9 mutagenesis of tomato *ADC* and *ODC* genes

Previously created tomato single *adc1* and *adc2* and double *adc1/adc2* mutants (Wu et al., 2019) and the new *odc1* mutant were generated via CRISPR-Cas9 (Jacobs et al., 2015) with two gRNAs, and transformed into tomato (Wittmann et al., 2016). Genotyping was conducted as described by Wu et al., 2019 and in Supplementary Figure 8.

### Phenotyping tomatoes

Germinated seedlings were genotyped, then transferred into 11 cm diameter pots at 18 DAG, and phenotypes measured/counted every 3 days from 26-47 DAG. Plant height was measured to nearest 0.5 cm from the cotyledon to shoot apical meristem. Only fully expanded adult leaves originated from the primary stem were counted. All buds and opened flowers were counted from each plant. Photos of every replicate were taken 37 DAG and leaf and flower photos taken at 47 DAG.

### Cold treatment of tomato leaves

The youngest, fully expanded adult leaf was cut from each replicate (aged 4-5 weeks). The terminal and two primary leaflets were kept in water at ∼6 °C or RT for 24 hours (in darkness). Samples for LC-MS or RNA extraction were taken without allowing CT leaves to warm.

### Generation of expression constructs

Tomato *ADC1*, *ADC2*, and *ODC1* genes were cloned from WT tomato gDNA into pUC57 vectors as level I modules for further Golden Gate cloning (Binder et al., 2014). Site-directed mutagenesis of key SlADC catalytic residues was conducted in level I modules. Constructs were cloned into T-DNA expression vectors and transformed into *A. tumefaciens GV3101*.

### Transient expression in *N. benthamiana*

*A. tumefaciens* strains containing T-DNA expression constructs and a p19 silencing suppressor plasmid-containing strain were grown in YEB media (28 °C, 180 rpm), centrifuged, and resuspended in infiltration buffer (10 mM MgCl_2_, 5 mM MES, and 150 µM acetosyringone) to an OD_600_ of ∼0.5. After an additional incubation (∼4 hours, 28 °C, 180 rpm), all expression construct-containing cultures were mixed 1:1 with the p19-containing strain to a final OD_600_ of 0.4 and infiltrated into *N. benthamiana* leaves using a blunt end syringe.

### ADC and ODC activity assay sample preparation

50 mg of plant tissue was flash frozen, ground in pre-cooled adapters and homogenized in ADC activity buffer (300 µL of 5 mM Tris [pH 7.5] with 0.75% polyvinylpolypyrrolidone [PVPP], 1 mM ascorbate, 50 µM P5P, 0.5x protease inhibitor [Roche complete], 2.5 µM D_5_ tryptophan [Sigma-Aldrich®] and 2µM ^13^C_1_D_2_ citrulline [Eurisotop®] as internal standards, 500 µM ^13^C_6_ arginine [Carl Roth®; for ADC activity], and 500 µM ^13^C_5_^15^N_2_ ornithine [Eurisotop®; for ODC activity]), keeping tubes on ice as much as possible. When testing for LDC activity, 500 µM ^13^C_6_^15^N_2_ lysine (LGC Standards®) was included in the buffer. Samples were incubated (1 hour, 37 °C, 350 rpm), then transferred to ice, and all reactions stopped with 5 µL concentrated HFBA (Sigma-Aldrich®), before centrifugation (18,000 g, 30 minutes, 4 °C). For *N. benthamiana* samples, 80 µL of the stopped assay samples was combined with 200 µL methanol and 100 µL chloroform, and incubated (10 minutes, 25 °C, 950 rpm). 120 µL of MilliQ water was added, the incubation repeated, followed by a final centrifugation (18,000 g, 30 minutes, 4 ° C), with the SN used for profiling.

### LC-MS profiling

Dilutions (in water, 0.1% FA, 0.1% HFBA) of representative samples were tested if their metabolite response fell into the linear range of the respective calibration curves (Supplementary Table 1). LC-MS profiling was performed using a Micro-LC M5 (Trap and Elute) and QTRAP6500+ (Sciex) system operated in MRM mode. Chromatographic separation was achieved on a HaloFused C18 micro column (150 x 0.5 mm; 2.7 μm; AB Sciex) and a Luna C18(2) micro trap column (5 μm; 100 Å; 20×0.5 mm; Phenomenex); injection volume was 50 μL. Analytes were ionized using an Optiflow Turbo V ion source equipped with a SteadySpray T micro electrode (10-50 μL min^-1^) in positive ion mode. Supplementary Table 2 presents information concerning the chromatographic gradients and MS parameters. A shorter trap gradient was chosen for the assays without HFBA in the solvent to reduce the loss of the ion paring agent in the sample itself. A general loss of sample volume of about 10% (data not shown) using the reduced trapping time was deemed acceptable.

Vendors Sciex OS software was used to analyze acquired MRM data. The metabolite content in each replicate was calculated using commercial standards (external calibration in water, 0.1% FA, 0.1% HFBA), except for ^13^C_5_ agmatine, ^13^C_4_ putrescine, and ^13^C_4_^15^N_2_ putrescine, which were quantified based on their natural variants. Data were normalized against sample FW, and D_5_ tryptophane or ^13^C_1_D_2_ citrulline.

*Note:* A small ^13^C_5_ agmatine peak was observed in control assays with no plant extracts. As no enzyme was present in these samples, the production of ^13^C_5_ agmatine can only be explained by chemical decomposition or insource fragmentation of ^13^C_6_ arginine within the MS ion source. This is further confirmed by the retention time of this ^13^C_5_ agmatine peak, which is not congruent to the natural agmatine but rather to arginine. All samples with low ADC activity (e.g., tomato *adc1/adc2* double mutant extracts) were double-checked for accumulation of enzymatically produced ^13^C_5_ agmatine (retention time of agmatine) instead of the chemically derived variant (retention time of arginine) (Supplementary Figure 10).

### RT-qPCR

RNA was extracted from flash frozen tomato samples using the Universal RNA Purification Kit (Roboklon GmbH); on-and oc-column DNase (ThermoFischer Scientific) treatments were both conducted as tomato *ADC*s do not contain introns. RNA (850 ng) was converted to cDNA using the RevertAid cDNA Synthesis Kit (ThermoFischer Scientific) and diluted 1:10 in RNA-free water. RT-qPCRs were conducted as described by Wu et al., 2019, using the CFX96 system (BioRad), and 3 technical replicates per biological replicate. *SlADC1* and *SlADC2* transcript levels were calculated using 2^-ΛλΛλCT^ and normalized to *SlTIP41* (reference gene) expression.

### Immunodetection of epitope-tagged proteins

*N. benthamiana* leaf tissue was flash frozen and ground in pre-cooled adapters, then homogenized in extraction buffer (80 mM Tris [pH 7.5], 2.5% SDS, 50 mM EDTA, 2.5x strength protease inhibitor; 33 µL per leaf disk) and loading dye added (20 mM Tris-HCl pH 8.6, 40% [v/v] glycerol, 10% [v/v] 2-mercaptoethanol, and 0.1% [w/v] bromophenol blue; 13 µL per leaf disk). Samples were heated (90 °C, 10 minutes), briefly centrifuged, and loaded onto a 10 % SDS PAGE gel. Proteins were transferred to a PVDF membrane using Trans-Blot® Turbo Transfer System (BioRad), blocked, then incubated overnight (6°C, rocking) with the epitope-detecting antibody (i.e., monoclonal, conjugated anti-HA-HRP antibody [Roche] or monoclonal anti-c myc [raised in mouse; Sigma Aldrich]). Washed blots (3x 5 min in TBST) were either imaged directly or blocked, incubated with a conjugated anti-mouse-HRP antibody (Sigma Aldrich) for 2 hours, and washed again. ECL Select^TM^ Western Blotting Detection Reagent (Amersham^TM^, cytiva) was used for detection (Amersham^TM^ Imager 600, GE Healthcare Life Sciences). Ponceau staining (0.1% [w/v] Ponceau S, and 5% [v/v] glacial acetic acid) was conducted to verify even protein loading.

### Data analysis and visualization

Microsoft Excel and R were used for basic statistical analysis and generation of graphs and heatmap diagrams, which were then assembled using Adobe Illustrator. Code to generate metabolite heatmaps is available: https://github.com/TheDepe/Metabolite-Measurement.

## RESULTS

### OPTIMIZATING THE LC-MS-BASED METHOD FOR ADC ACTIVITY QUANTIFICATION

We first sought to improve the sensitivity of our LC-MS-based method to quantify ADC activity and PA-related metabolites from plant extracts, even with low *ADC* expression. In the previously established protocol, ADC activity was quantified in the supernatant (SN) of cellular extracts (Figure 2B; Wu et al., 2019). We rationalized that SNs may lack organelles containing ADC proteins and metabolites (e.g., chloroplasts), consequently reducing ADC activity. Therefore, we compared ADC activity in SNs (i.e., previous protocol) to whole cellular extracts (termed here on out as ‘whole extract’ or WE; Figure 2B). Furthermore, we considered whether a higher ADC substrate (i.e., ^13^C_6_ arginine; Figure 2A) concentration would elevate ADC activity. We therefore compared using ten times more ^13^C_6_ arginine (i.e., 500 µM; “high”) to the original 50 µM (“low”) concentration (Figure 2B).

High versus low concentrations of the ADC substrate were added to the SN or WE of tomato leaves kept at either room temperature (RT) or cold-treated (CT); the latter is known to elevate *SlADC* transcript levels (Upadhyay et al., 2020). Indeed, both *SlADC1* and *SlADC2* transcript levels were significantly induced by ∼3.5- and 2.9-fold following CT relative to RT controls (Supplementary Figure 1). When using the higher ^13^C_6_ arginine concentration, we observed a dramatic increase in ADC activity (Figure 2C). While this was particularly evident in CT samples with induced *SlADC* levels (∼11-fold for WE and ∼7-fold for SN samples), a similar increase was observed in RT samples (i.e., ∼11- and ∼8-fold higher in WE and SN samples, respectively), although not statistically significant (Figure 2C). Interestingly, applying a higher ^13^C_6_ arginine concentration enabled reliable detection of the heavy isotopic variant of putrescine produced from ^13^C_6_ arginine (i.e., ^13^C_4_ putrescine; Figure 2A), which was not detectable with the low ^13^C_6_ arginine concentration (Figure 2D). Overall, these results demonstrate that increasing the ^13^C_6_ arginine substrate concentration elevated the sensitivity of our ADC activity assay, enabling reliable measurement of ^13^C_5_ agmatine, and derived ^13^C_4_ putrescine, even under *ADC* non-inducing conditions.

Unexpectedly, we observed no significant differences in levels of ^13^C_5_ agmatine or ^13^C_4_ putrescine between SN and WE samples (Figure 2C and D). However, the SN-based ADC activity protocol is more labor-intensive, time-consuming, and error-prone due to more handling steps, making the new WE protocol preferable.

### ADDITION OF ODC ACTIVITY QUANTIFICATION TO OUR LC-MS METHOD

Measuring multiple enzyme activities in parallel reduces experimental costs and workload. Therefore, we wanted to test if both ADC and ODC activities (i.e., decarboxylation of ornithine to putrescine) could be quantified in one LC-MS run, despite both pathways producing putrescine (Figure 1). To achieve this, heavy isotopically labelled substrates of ADC and ODC were selected that would produce putrescine variants with sufficiently different molecular masses to be distinguishable by MS. For this, ^13^C_6_ arginine (+6 mass) and ^13^C_5_^15^N_2_ ornithine (+7 mass) were selected as ADC and ODC substrates, which produce ^13^C_4_ putrescine (+4) and ^13^C_4_^15^N_2_ putrescine (+6), respectively (Figure 2A and E).

We successfully detected ^13^C_4_^15^N_2_ putrescine in our tomato extracts and discriminated it from the ADC-derived ^13^C_4_ putrescine, enabling simultaneous ADC and ODC activity quantification (Figure 2F). In contrast to ADC activity, ODC activity was not elevated by higher substrate levels and ODC activity was higher in all WE assays compared to the SN. Our results indicate that a substantial portion of ODC enzymes are localized in organelles and suggest that WEs are preferable to SNs for assaying enzyme activities.

### THE ION PAIRING AGENT HFBA, IS NOT AN ESSENTIAL COMPONENT OF THE LC-MS GRADIENT COMPOSITION

We also considered the LC-MS gradient composition for optimization. Previously, 0.05% heptafluorobutyric acid (HFBA) was included in the gradient solvents as an ion pairing agent (Wu et al., 2019, Gallas et al., 2024). However, due to its acidity, HFBA could cause corrosion of internal LC-MS components and also increase LC-MS contamination levels by adhering to the column and MS ion source. This prevents quick switching between positive and negative ionization modes, as its acidiff nature creates a substantial matrix suppression effect (Jessome and Volmer, 2006). The assay from Figure 2 was analyzed twice; once with HFBA in the solvent, and once without, relying solely on the HFBA added to the sample (i.e., to stop enzyme activity assay; Figure 2B) for ion pairing. In all cases, ^13^C_5_ agmatine intensities showed no significant difference between the two gradient compositions (Supplementary Figure 2). Therefore, HFBA can be omitted from the solvent without affecting activity quantification, whilst improving longevity and performance of the LC-MS system.

### EXPANDING THE PROFILE OF QUANTIFIABLE PA-RELATED METABOLITES ENABLES INTERPRETATION OF METABOLIC FLUXES AFTER COLD TREATMENT

Monitoring changes across the wider PA network – particularly of anabolic compounds – would improve our understanding of PA metabolic fluxes, especially given the known shifts in amino acid levels during stress responses (Batista-Silva et al., 2019, Paschalidis et al., 2019, Heinemann and Hildebrandt, 2021, Moormann et al., 2022). Previously, we had only quantified the concentrations of agmatine, putrescine, spermidine, and spermine (Wu et al., 2019, Gallas et al., 2024). Yet, to better comprehend the impact of abiotic stresses or extraction methods (e.g., SN and WE), we extended our protocol to include arginine, ornithine, citrulline, glutamine, N-acetylputrescine, cadaverine, and later, lysine. This was achieved by adjusting LC gradients to avoid co-elution of structural isomers such as acetyl-putrescine and agmatine, and glutamine and lysine (Supplementary Table 1 and 2, Supplementary Figure 3).

When quantifying changes in the concentrations of numerous metabolites in response to external (e.g. biotic or abiotic stress) or internal (e.g. developmental program) stimuli, or due to differences in extraction protocols (e.g., SN versus WE), it is difficult to infer metabolic fluxes if the data are displayed using conventional bar charts, as such representations do not reveal the causal relationships between metabolites. Therefore, we presented our LC-MS profiling results as a heatmap-based pathway diagram to better visualize metabolic relationships of the PA network (Figure 3). Following CT, agmatine, putrescine, ornithine, and citrulline levels increased, while arginine levels were reduced compared to the RT samples. This suggests that cold-induced *SlADC* transcript levels and ADC activity reduces the arginine pool to produce more agmatine and subsequently putrescine. Interestingly, spermidine and spermine levels are not influenced by CT, suggesting a regulatory mechanism that prevents the cold-induced increase in putrescine from causing an unregulated increase in these higher PAs.

**Figure 3:**
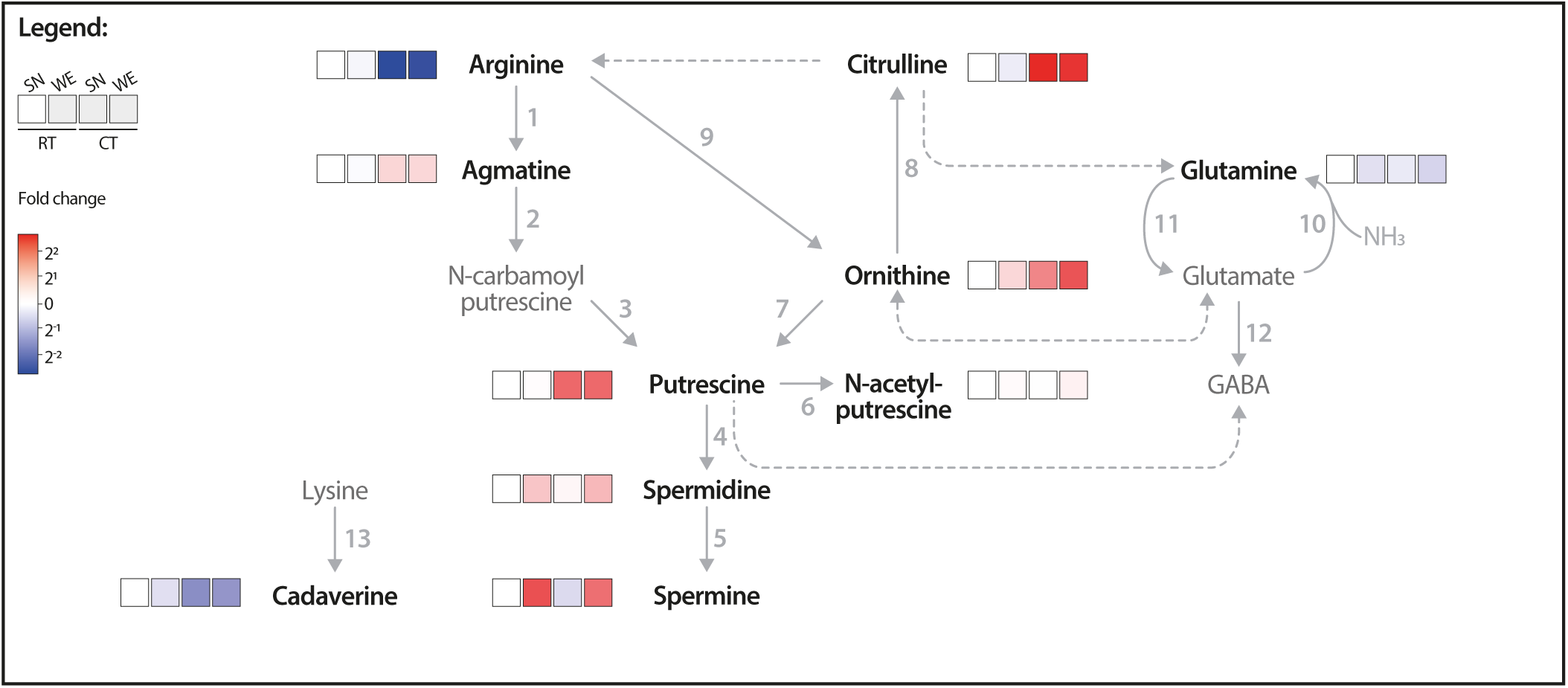
Pathway heatmap highlights how cold treatment influences PA network flux. PA and related metabolite concentrations were measured via LC MS from room temperature (RT) and cold treated (CT) tomato leaf samples from either supernatant (SN) or whole extraction (WE) assay samples. Only samples with 500 µM 13C6 arginine used are shown. The log2 fold change (FC) of each metabolite was calculated relative to the RT, SN-extracted (RT SN) condition; red indicates higher and blue lower metabolite levels compared to the RT SN. Solid arrows represent reactions catalyzed by a single enzyme, while dotted arrows indicate multiple enzymatic steps. Numbers correlate to the following enzymes: 1, ADC; 2, AIH; 3, NCPAH; 4, SPDS; 5, SPMS; 6, NATA1; 7, ODC; 8, OTC; 9, ARGAH; 10 GOGAT; 11, GS; 12, GAD; and 13, LDC. Only bold, black named metabolites were measured in this LC MS experiment.

Our results collectively show that a higher substrate concentration (i.e., 500 µM) increases assay sensitivity. Additionally, using WEs instead of SNs, now captures the activity of both cytoplasmic- and organelle-localized enzymes. We also found that omitting the ion pairing agent HFBA from the LC-MS gradient had no impact metabolite quantification but protects the LC-MS mechanical components. Finally, visualizing PA-related metabolite changes via a heatmap pathway diagram aids in understanding the interdependencies between metabolites.

### ADC PROTEIN ABUNDANCE CORRELATES WITH ADC ACTIVITY

Previously, we assumed a linear relationship between ADC protein abundance and its enzymatic activity; yet this assumption had not been experimentally validated (Wu et al., 2019, Gallas et al., 2024). To test for such a linear relationship, we expressed tomato ADC (SlADC) proteins in *N. benthamiana* leaves, where their expression levels can be easily modulated by inoculating varying amounts of T-DNA-containing *A. tumefaciens* strains. These T-DNA constructs encoded epitope-tagged SlADC1/2, driven by the constitutively active *35S* promoter. They were transformed into *A. tumefaciens* and infiltrated into *N. benthamiana* leaves (termed agroinfiltration) with increasing bacterial loads (i.e., OD_600_ 0.05, 0.1, 0.5, and 1). Two days post infiltration (dpi), samples were harvested for both immunoblotting and LC-MS profiling. Immunoblots showed increasing signal intensity of tagged SlADCs that mirrored the increasing bacterial loads (Figure 4). Previously, unprocessed *N. benthamiana* leaf samples rapidly clogged the micro-LC columns, probably due to high alkaloid levels in this species (Stephan et al., 2018). Therefore, an additional liquid-liquid extraction step was applied to all *N. benthamiana* leaf samples following the assay. Comparable to the protein immunodetection analysis, rising bacterial load increased ADC activity (Figure 4), confirming our hypothesis of a linear relationship between ADC abundance and activity. To confirm that this observation was not due to heightened biotic stress influencing ADC activity, the same experiment was conducted with epitope-tagged fluorophore controls. Again, increasing bacterial loads were associated with higher detectable epitope-tagged protein levels, but no mirrored increase in ADC activity was observed (Supplementary Figure 4). Overall, these results confirm a linear relationship between SlADC1/2 protein abundance and ADC activity, demonstrating that ADC activity quantification can be used as a reliable proxy for ADC protein levels from plant extracts.

**Figure 4:**
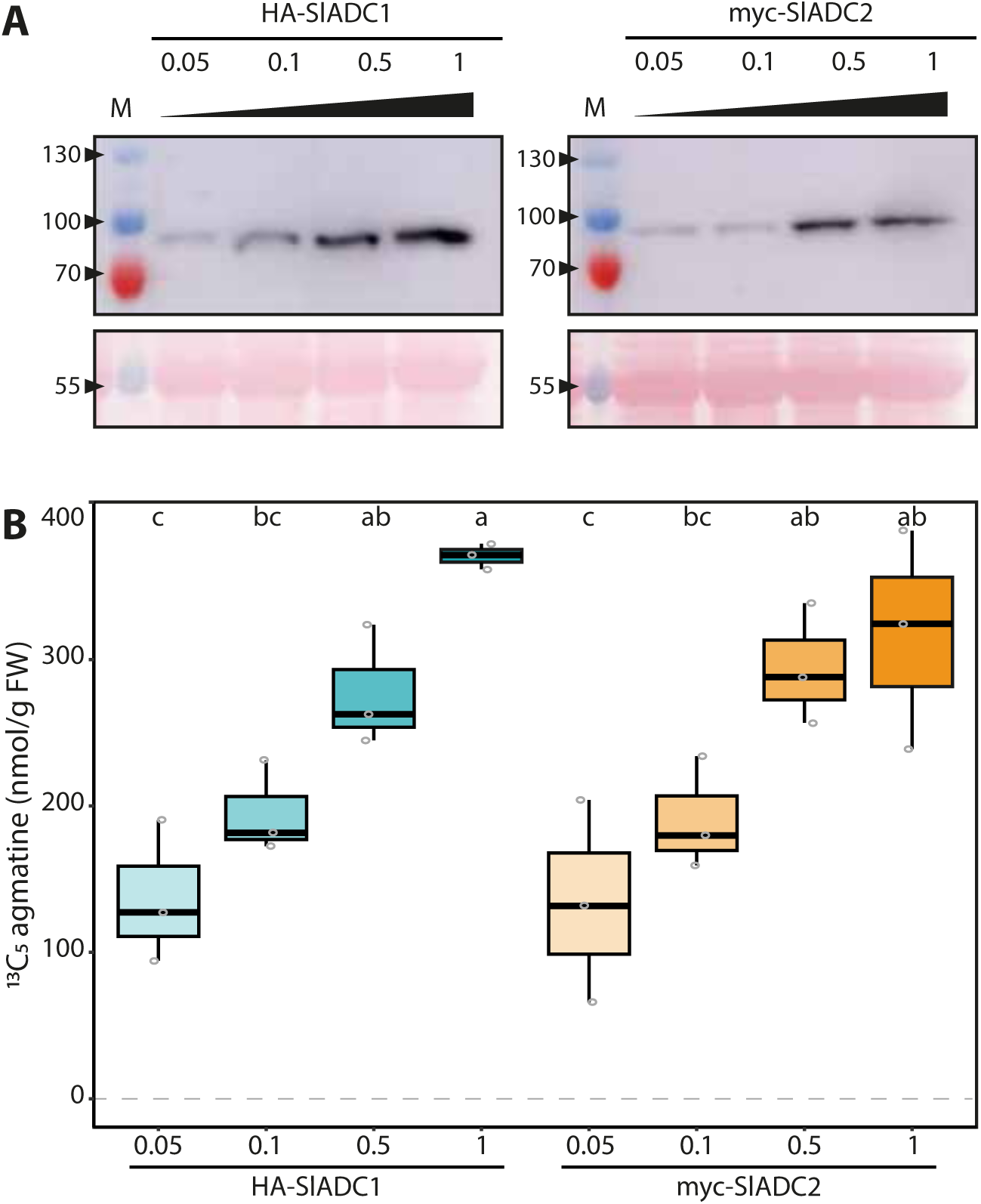
ADC activity directly and linearly correlates with ADC protein abundance. **A)** ADC1 and ADC2 protein abundance increases with increasing bacterial load. Immunoblot analysis of *N. benthamiana* leaves agroinfiltrated with 35Sp:HA-SlADC1:35St (left) and 35Sp:myc-SlADC2:35St (right) at increasing OD600 (0.05, 0.1, 0.5, and 1) of samples taken 2 days post infiltration (dpi). Upper images (cropped) show HA-SlADC1 detection (left) and myc-SlADC2 (right), expected sizes are 78 and 80 kDa, respectively. Ponceau stained blots show equal protein loading (bottom). **B)** ADC activity directly correlates with increasing ADC protein abundance. ADC activity (13C5 agmatine in nmol/g FW, normalized to D5 tryptophan) from the same *N. benthamiana* HA-SlADC1 (blue) and myc-SlADC2 (orange)-agroinfiltrated samples shown in A), with increasing color intensity representing increasing bacterial loads. Statistical significance was determined using an ANOVA followed by Tukey’s post hoc HSD test; different letters above each box indicate statistically significantly different groups (a = 0.05). Each biological replicate represented by a grey circle; n = 3.

### ADC ACTIVITY QUANTIFICATION TO TEST FUNCTIONALITY OF WT AND MUTANT ADC VARIANTS

We next wanted to evaluate whether the enzymatic functionality of ADC mutant variants, recombinantly expressed in *N. benthamiana* leaves, can be assessed using our optimized ADC activity assay. Previously, Hanfrey et al., 2001 showed that *A. thaliana* ADC1 residues K136 and C524 were essential for enzymatic function. Using a peptide sequence alignment, we determined that these residues correspond to K149/K156 and C540/C548 in SlADC1 and SlADC2, respectively (Supplementary Figure 5). We mutated these residues either individually (i.e., SlADC1 K149A or C540A, and SlADC2 K156A or C548A) or both to alanine (i.e., SlADC1 K149A C540A and SlADC2 K156A C548A). T-DNA constructs were generated encoding *35S*-promoter driven, epitope-tagged versions of mutated *SlADC1/2*, transformed into *A. tumefaciens*, and agroinfiltrated into *N. benthamiana* leaves alongside complementary mCherry and GFP fluorophores, serving as negative controls. LC-MS analysis of wild-type (WT) SlADC1- and SlADC2-agroinfiltrated samples showed 72- and 85-fold higher ADC activity compared to the average ADC activity of the negative controls (Figure 5). No significant difference between the negative controls and any single or double SlADC1 or SlADC2 mutant variant was found, despite all proteins being equally expressed (Supplementary Figure 6). These results demonstrate that both lysine and cysteine residues are essential for SlADC1/2 catalytic activity.

**Figure 5:**
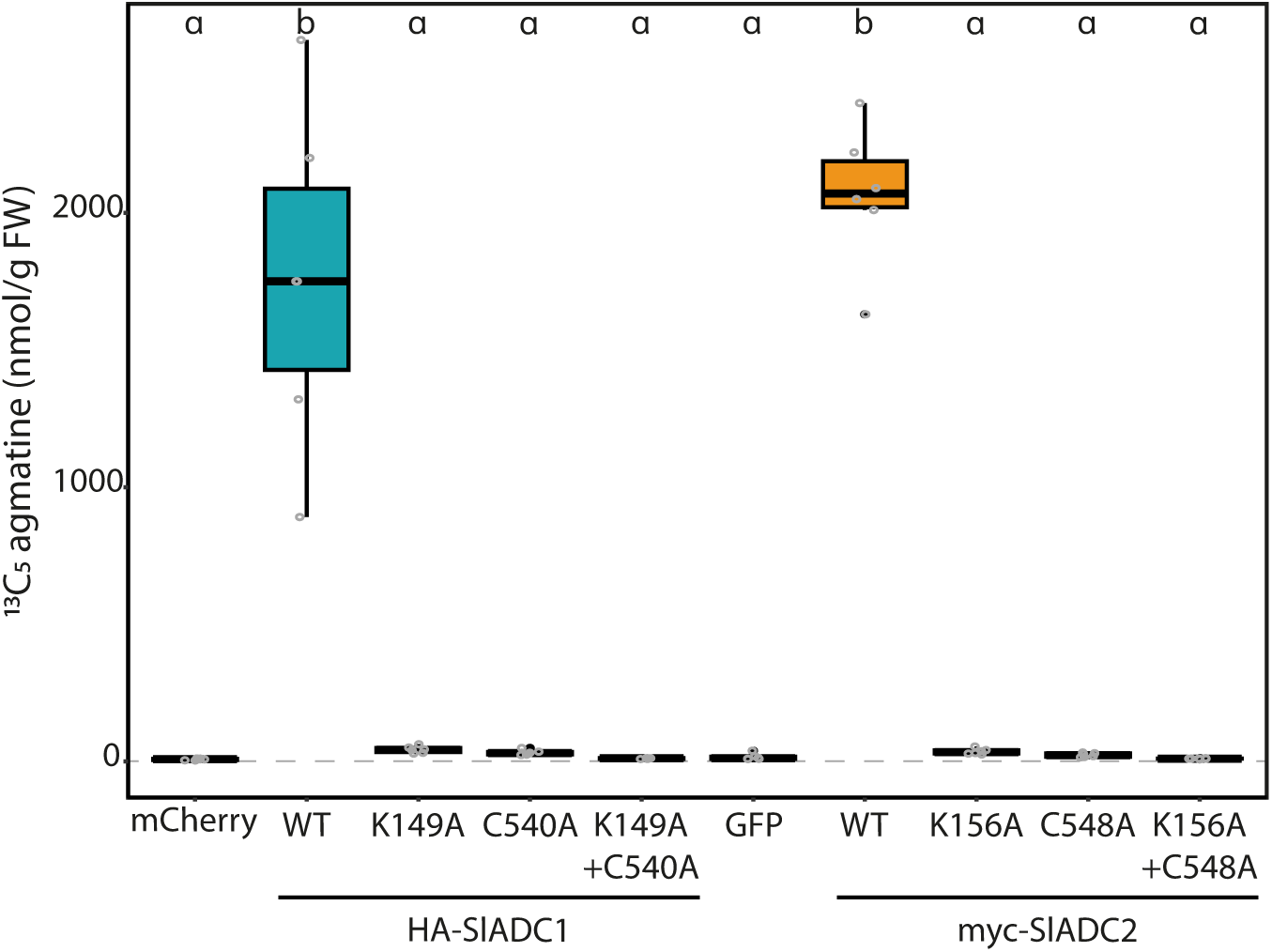
Mutating either the cofactor- or substrate-binding residues kills catalytic functionality of SlADC1 and SlADC2. **A)** *N. benthamiana* leaves were agroinfiltrated with 35S-promoter driven HA-SlADC1 (blue) or myc-SlADC2 (orange) WT or mutant variants, alongside HA-mCherry and myc-GFP fluorophores as negative controls. After 2dpi, ADC activity (13C5 agmatine in nmol/g FW, normalized to D5 tryptophan) was quantified using the optimized LC-MS method. Statistical significance determined using ANOVA followed by Tukey’s post hoc HSD test; different letters above each box indicate statistically significantly different groups (a = 0.05). Each biological replicate represented by a grey circle; n = 6.

### ANALYZING SUBSTRATE SPECIFICITY OF TRANSIENTLY EXPRESSED ADC AND ODC ENZYMES IN *N. BENTHAMIANA*

As ADCs and ODCs are key enzymes in PA biosynthesis, we wanted to show that our LC-MS assay could quantify both activities following their transient expression in *N. benthamiana*. We generated *35S*-promoter driven T-DNAs encoding, HA epitope-tagged SlADC1, SlADC2, SlODC1, and mCherry (negative control) and agroinfiltrated these into *N. benthamiana* leaves. At 2 dpi, samples were harvested for both LC-MS and immunoblot analysis. Immunoblot analysis showed that all proteins were detectable (Supplementary Figure 7).

Previous studies have suggested that some ODCs have promiscuous activities and will also decarboxylate lysine into cadaverine (Bunsupa et al., 2012). To determine substrate specificity, a ^13^C_6_^15^N_2_ lysine substrate was included in the activity assays along with the isotopic variants of arginine and ornithine.

As anticipated, transient expression of SlADC1 and SlADC2 produced 83- and 58-fold higher ADC activity levels compared to the negative mCherry control, whilst SlODC1 expression produced no significant increase in ADC activity (Figure 6A). This pattern was mirrored in the accumulation of ^13^C_4_ putrescine (generated from ^13^C_6_ arginine), where SlADC1 and SlADC2 expression caused 42- and 43-fold higher levels compared to the control (Figure 6B). On the other hand, 20-fold higher ODC activity was observed upon transient SlODC1 expression, whilst no significant increase was observed following SlADC1 and SlADC2 expression (Figure 6C). Interestingly, expression of SlADC1 and SlADC2 enzymes, but not SlODC1, significantly increased putrescine compared to the mCherry control (Figure 6D). Finally, no LDC activity (i.e., production of ^13^C_5_^15^N_2_ cadaverine from ^13^C_6_^15^N_2_ lysine) was detected above background levels from any expressed enzyme.

**Figure 6:**
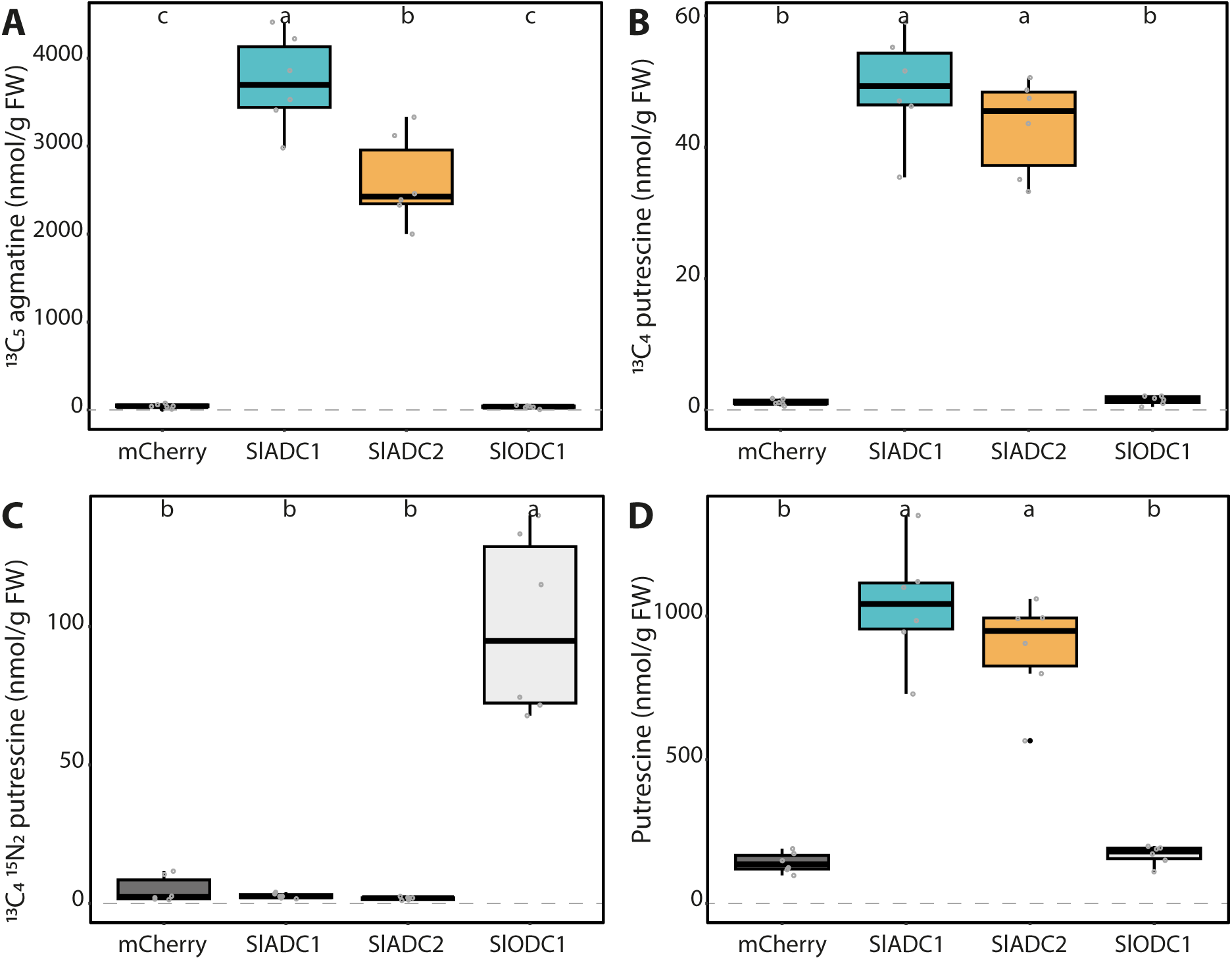
Transiently expressed tomato ADC and ODC enzymes show high substrate specificity. **A)** *N. benthamiana* leaves were agroinfiltrated with 35S-promoter driven HA epitope-tagged mCherry (dark grey), SlADC1 (blue), SlADC2 (orange), or SlODC1 (light grey) constructs. **A)** ADC activity (13C5 agmatine) and **B)** 13C4 putrescine can be detected following expression of tomato ADC enzymes. **C)** Only expression of tomato ODC1 produced high ODC activity (13C415N2 putrescine). **D)** Expression of tomato ADCs, but not ODC1, induced putrescine levels. All metabolite levels are presented in nmol/g FW, normalized to D5 tryptophan. Statistical significance determined using ANOVA followed by Tukey’s post hoc HSD test; different letters above each box indicate statistically significantly different groups (a = 0.05). Each biological replicate represented by a grey circle; n = 6.

Overall, these results demonstrate that tomato ADC and ODC enzymes have strict preference for their respective metabolic substrates and that their activities can be easily analyzed following transient expression in *N. benthamiana* leaves without the need for *in vitro* protein purification.

### CHARACTERIZATION OF THE PA NETWORK AND PHENOTYPES OF TOMATO ADC AND ODC KNOCK OUT MUTANTS

To gain deeper insight into the PA network, we analyzed tomato *adc1*, *adc2*, and *adc1/adc2* mutants (Wu et al., 2019) using our optimized LC-MS profiling method. We also included an unpublished single *odc1* tomato mutant, with the remaining two *SlODC* genes WT (*Solyc03g098300.1* and *Solyc03g098310.1*). These tomato lines were all generated via CRISPR-Cas9 mutagenesis and carry deletions that lead to premature stop codons (Wu et al., 2019; Supplementary Figure 8). In contrast to *A. thaliana* where the loss of both *ADC* genes is embryonically lethal due to the absence of *ODC*, the tomato *adc1/adc2* double mutant is viable, but cannot produce seeds. Since *adc1/adc2* mutants produce no seeds, we identified these double mutants by genotyping the progeny of a parental line that was homozygous mutant for *SlADC1* (sl*adc1/sladc1*) and heterozygous for *SlADC2* (*SlADC2/sladc2*) (Supplementary Figure 8D). Since *adc1/adc2* mutants offer a unique opportunity to study the functional roles of ADCs, we analyzed phenotypic differences between these mutants and WT plants. Twenty genotyped tomato plants representing WT, *adc1*, *adc2*, *adc1/adc2*, and *odc1* were examined every three days between 26 to 47 days after germination (DAG), with photos taken of representative plants at 37 DAG (Figure 7A). Leaves of *adc1/adc2* mutants developed primary and secondary leaflets more slowly, often lacking intercalary leaflets entirely (Figure 7B). By 47 DAG, their leaves and leaflets were less flat and expanded compared to those of WT plants. Interestingly, no flowers, buds, or tissues indicating developing flower buds were ever observed on *adc1/adc2* plants, demonstrating that *ADC* genes are indispensable for flower development (Figure 7C and D). By contrast, the single *adc1* or *adc2* mutants showed no obvious deficiency in flower development compared to WT. Additionally, the double *adc1/adc2* mutant plants were shorter and had fewer adult leaves compared to WT or the other mutants (Figures 7E and F). Intriguingly, *adc1* and *odc1* were both significantly taller 47 DAG compared to the WT. Overall, this phenotypic analysis demonstrates the importance of ADCs for tomato growth and development of reproductive organs.

**Figure 7:**
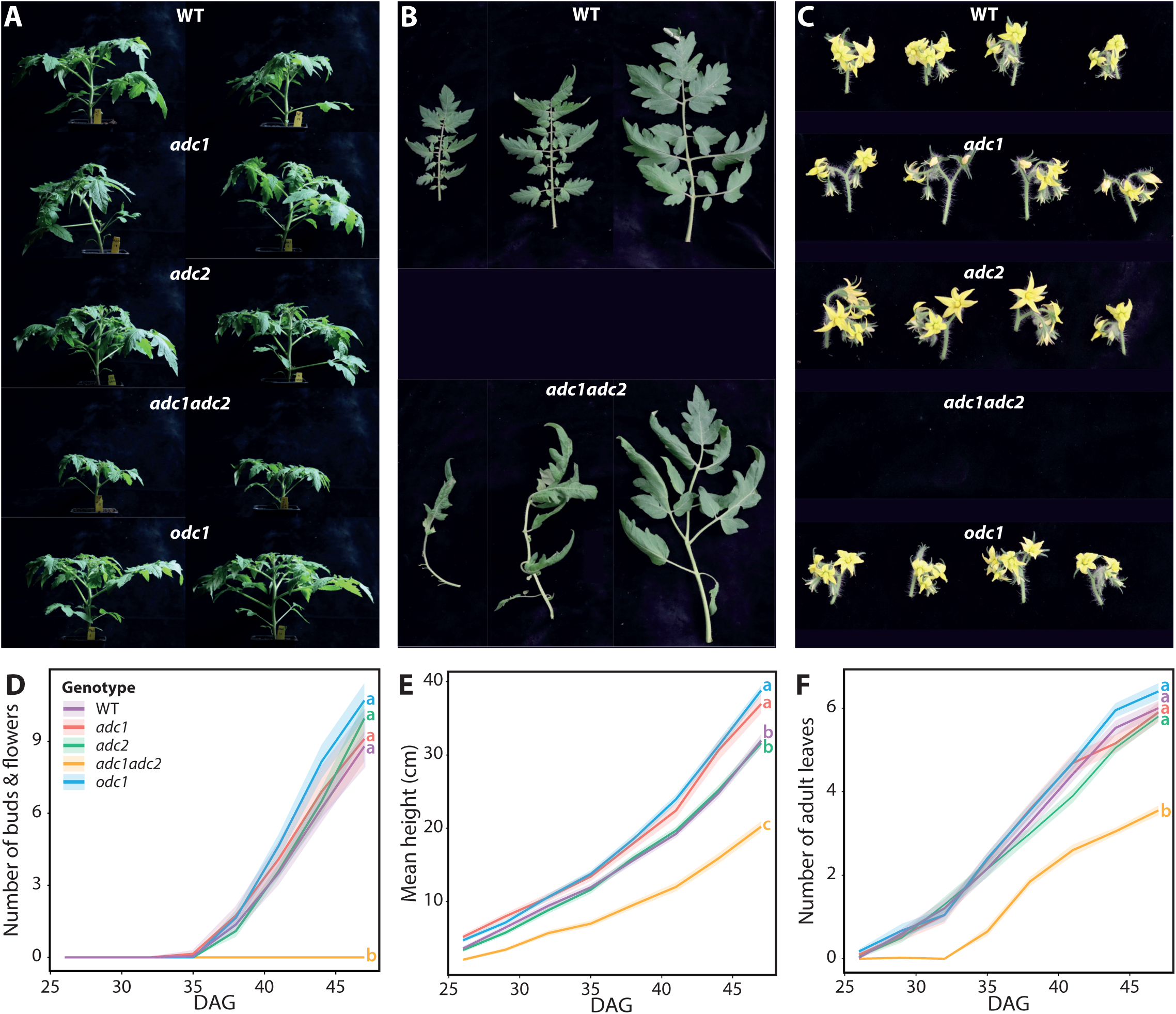
Tomato *adc1/adc2* mutant is smaller with fewer leaves and inability to form flowers. Phenotypes of WT, *adc1*, *adc2*, *adc1/adc2*, and *odc1* tomato mutants were analyzed until 47 days after germination (DAG). **A)** Photos of whole plants were taken at 37 DAG; two representatives are shown. **B)** Leaves of the *adc1/adc2* double mutant have morphological differences to WT. Photos taken 47 DAG. **C)** At 47 DAG, photos were taken of flowers from each genotype, except for *adc1/adc2* as no flowers were ever formed. **D)** Flower and bud numbers were counted. **E)** Double *adc1/adc2* mutant plants were also shorter and **F)** had fewer adult leaves. All plants (WT [purple], *adc1* [red], *adc2* [green], *adc1/adc2* [yellow], and *odc1* [blue]) were analyzed every 3 days from 26-47 DAG; n = 20. Solid lines represent mean and shaded area represents standard error of the mean. Statistical significance was determined only on data from 47 DAG; an ANOVA followed by Tukey’s post hoc HSD test was conducted; different letters above each box indicate statistically significantly different groups (a = 0.05).

To determine the role each ADC enzyme plays in PA metabolism, transcript levels, ADC activity and PA-related metabolites were quantified in all five genotypes following an acute CT. *SlADC1* and *SlADC2* transcript levels were increased in all genotypes following CT, except for *SlADC1* in the *adc1* mutant (Supplementary Figure 9). ADC activity was detectable for all genotypes following RT or CT, except for the *adc1/adc2* double mutant (Figure 8A), consistent with agmatine being exclusively synthesized by ADCs. Following CT, ADC activity was significantly induced in the WT, *adc2*, and *odc1* genotypes, but not *adc1*, despite significant transcriptional activation of its functional *SlADC2* gene following CT (Supplementary Figure 9B). These results indicate that SlADC1 is the main contributor to ADC activity in adult tomato leaves following CT.

**Figure 8:**
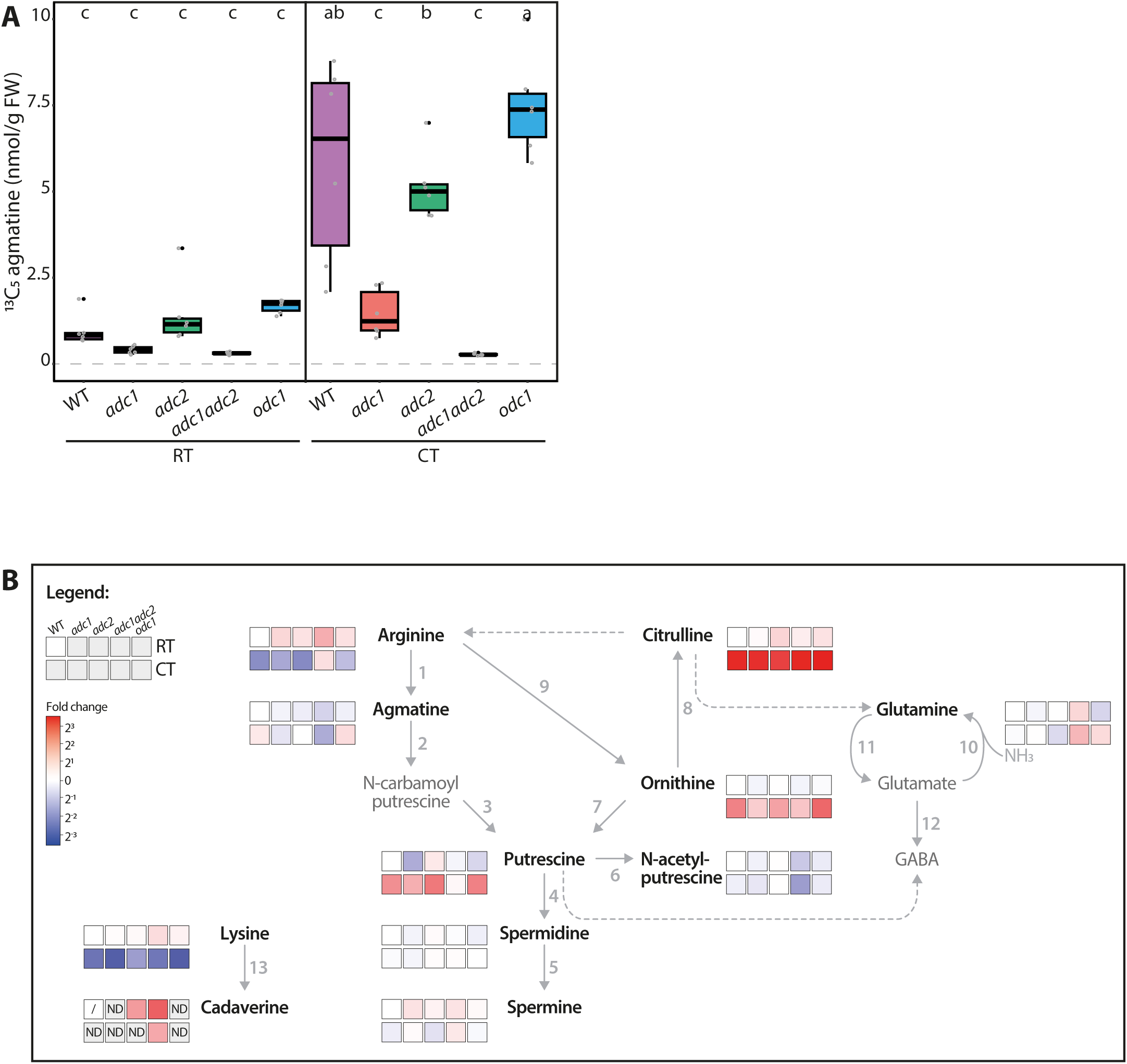
Tomato adc/1adc2 mutant has no ADC activity and perturbed PA metabolism. **A)** ADC activity of *adc1* and *adc1/adc2* is severely affected in both room temperature (RT) and cold treated (CT) conditions. ADC activity (13C5 agmatine in nmol/g FW, normalized to D5 tryptophan) of WT (purple), *adc1* (red), *adc2* (green), *adc1/adc2* (yellow), and odc1 (blue) adult tomato leaves following RT or CT at 38 DAG. Each biological replicate represented by a grey circle (n = 6). Statistical significance determined using ANOVA followed by Tukey’s post hoc HSD test; different letters above each box indicate statistically significantly different groups (a = 0.05). **B)** Heatmap pathway representation demonstrates impact of knocking out ADCs on PA metabolism in tomatoes. Prior to the assay incubation, 100 µL of sample was removed and immediately stopped with HFBA; this was analyzed by LC MS. The log2 fold change (FC) of each metabolite was calculated relative to the RT-treated WT (WT RT) control; red indicates higher and blue lower metabolite levels compared to the WT RT. Solid arrows represent reactions catalyzed by a single enzyme, while dotted arrows indicate multiple enzymatic steps. If a metabolite was not detected (ND) the box is shaded grey. A metabolite not detected in all biological replicates of the WT RT reference is symbolized with a dash. Numbers correlate to the following enzymes: 1, ADC; 2, AIH; 3, NCPAH; 4, SPDS; 5, SPMS; 6, NATA1; 7, ODC; 8, OTC; 9, ARGAH; 10 GOGAT; 11, GS; 12, GAD; and 13, LDC. Only bold, black named metabolites were measured in this LC-MS experiment.

When quantifying PA metabolites, we again observed higher agmatine, putrescine, citrulline, and ornithine levels and lower arginine levels in WT tomato leaves following CT (Figure 8B). Contrastingly, the *adc1/adc2* double mutant had no detectable agmatine above background levels and under both RT and CT conditions the *adc1/adc2* mutant had higher arginine but lower putrescine levels. Interestingly, N-acetylputrescine levels were also lower in the *adc1/adc2* mutant compared to WT, while spermidine and spermine levels remained unaffected. Finally, we noted that although lysine (the cadaverine precursor) was lower in all genotypes following CT, cadaverine was not increased and was only reliably detected in the *adc1/adc2* mutant under both conditions.

In summary, we characterized the phenotypes, ADC activities, and PA network in the tomato *adc1*, *adc2*, *adc1/adc2*, and *odc1* mutants, demonstrating the importance of ADC enzymes not only for PA synthesis, but also for plant development and generation of reproductive tissues.

## DISCUSSION

### Optimizations vastly improved the previous LC**-**MS-based ADC activity assay

In this work we optimized the sensitivity and workflow of our previously established LC-MS-based method for quantifying ADC activity and PAs from plant extracts. Moreover, we broadened the scope of this method, including ODC activity and 7 more PA-related metabolites (i.e., arginine, ornithine, citrulline, glutamine, N-acetylputrescine, lysine, and cadaverine) that were not covered by the previous method. Multiple aspects of our LC-MS approach were considered for optimization, including substrate concentration, extraction method, and presence of the ion-pairing agent HFBA in the solvent.

#### Higher substrate concentration increases ADC activity levels

Using 10-fold higher substrate concentration increased ADC activity ∼11-fold (Figure 2C), producing clear ^13^C_5_ agmatine LC-MS peaks well above detection limits, even from samples with low levels of ADC proteins (e.g., RT single *adc* mutants; Figure 8). The ^13^C_6_ arginine-derived putrescine variant (^13^C_4_ putrescine) could only be detected with elevated substrate levels, but not at lower ^13^C_6_ arginine concentrations (Figure 2D). Since at both substrate concentrations ^13^C_6_ arginine levels were never depleted (Supplementary Data File), the higher concentration (i.e., 500 µM) likely increased ADC activity by almost reaching the reported SlADC1 Km of 600 µM (Guo et al., 2023), a situation representing 50% substrate saturation of the enzyme. A higher ^13^C_6_ arginine concentration to achieve 100% enzyme substrate saturation was not considered as this could negatively affect the chromatographic separation and electrospray ionization process of coeluting metabolites (Jessome and Volmer, 2006). Taken together, our results suggest that 500 µM ^13^C_6_ arginine increased assay sensitivity, facilitating detection of subtle fluctuations in ADC activity that occur in response to abiotic stresses.

#### Simultaneous quantification of ADC and ODC activity from plant tissue

Putrescine, the precursor for synthesizing higher plant PAs, is produced via the ADC- or ODC-dependent pathways. We aimed to establish an approach allowing quantification of both ADC and ODC enzymatic activities from the same tissue extract in one LC-MS sample. To quantify ODC activity, we used a stable ^13^C_5_^15^N_2_ ornithine isotope that, following decarboxylation by ODC, produces ^13^C_4_^15^N_2_ putrescine. This putrescine isotope differs in its molecular mass sufficiently from the ADC-derived ^13^C_4_ putrescine (i.e., from ^13^C_6_ arginine), enabling quantification of both ADC and ODC activities simultaneously (Figure 2F). In contrast to ADC activity, increasing ^13^C_5_ ^15^N_2_ ornithine substrate concentration did not elevate ODC activity. This is likely due to low SlODC protein levels as *SlODC* is not highly transcribed in adult tomato leaves (Kwak and Lee, 2001, Acosta et al., 2005, Sivakumar et al., 2022).

#### ODC activity is higher in whole extracts containing all organelles

We compared ADC and ODC activities in WEs and SNs (Figure 2B) and consistently found higher ODC activity in WEs than SNs (Figure 2F). ODCs have been shown to localize to the endoplasmic reticulum (ER, Joshi et al., 2024) and predicted to localize to chloroplasts and mitochondria (Sivakumar et al., 2022). Centrifugation to obtain SNs removes these key organelles, thereby reducing ODC abundance and activity. In contrast, ADC activity did not differ between SNs and WEs (Figure 2C), indicating that SlADCs are predominantly localized in the cytoplasm of tomato leaf cells. Overall, this work demonstrates that analysis of WEs simplifies the workflow and is equally suitable for enzymes localized in the cytoplasm, ER or plastids. Comparative LC-MS analysis of SN and WE samples could also provide information regarding localization of natively expressed enzymes, thereby complementing classical subcellular localization studies.

#### Additional HFBA in the LC-MS gradient is not required for high quality metabolite quantification

Finally, we tested whether HFBA could be omitted from the LC-MS gradient whilst still enabling reliable ADC activity quantification. HFBA was used for two purposes in the previously published method: firstly, to stop proteinaceous reactions (i.e., enzymatic activity) at a defined timepoint (Figure 2B); and secondly, as an ion pairing agent, that improves resolution of charged metabolites. However, HFBA potentially reduces LC-MS platform longevity and integrity, and creates matrix suppression effects. Upon comparing metabolite peak areas of the same samples analysed with or without HFBA in the LC-MS gradient, we observed no significant differences (Supplementary Figure 2), indicating that sufficient ion-pairing occurs from the HFBA added directly into the samples. Therefore, HFBA can be omitted from the LC-MS gradient without compromising metabolite and enzyme quantification, improving LC-MS system longevity.

#### Studying post-transcriptional regulation: ADC and ODC activity quantification as a proxy for their protein abundance *in planta*

A gene’s transcript level is commonly used to estimate the amount and biological activity of the encoded protein, yet this approach is unsuitable for ADC and ODC, whose translation is also regulated post-transcriptionally (Kwak and Lee, 2001, Wu et al., 2019, Jimenez-Bremont et al., 2022). Here, we validated a linear relationship between ADC activity and ADC protein abundance (Figure 4), showing that enzyme activity can be used as a proxy for protein abundance *in planta*. Therefore, with the simplicity and reliability of our LC-MS approach, it is now possible to efficiently compare ADC and ODC activities, and accordingly their protein levels, across many tissues and/or stimuli.

Furthermore, systematic comparison of *ADC*/*ODC* mRNA levels with enzyme activities quantified using our LC-MS approach could identify contexts where transcript levels and enzyme activities differ most. As such contexts with large differences would be indicative of post-transcriptional regulation, our approach has the potential to be used in systematic screens, uncovering tissue contexts or stimuli where ADC and ODC expression is regulated at the post-transcriptional level.

#### Additional purification step opens the door for analysis of transiently expressed ADC and ODC enzymes

*Agrobacterium*-mediated transfer of T-DNAs encoding proteins of interest into *N. benthamiana* leaves has become a widely used method for functional analysis of plant genes. We therefore wanted to ensure that our LC-MS approach could be applied to ADC/ODC proteins ectopically expressed *N. benthamiana* leaves. Yet, when using the standard extraction protocol, these samples clogged our Micro-LC columns. By including an additional extraction step, *N. benthamiana* samples no longer interfered with the columns, enabling key experiments to be conducted (Figures 4, 5, and 6). Using the *N. benthamiana* expression system, we not only verified that ADC activity has a linear relationship to ADC protein abundance (Figure 4) but also confirmed the functional relevance of key lysine and cysteine residues required for SlADC activity (Figure 5). Compatibility of the *N. benthamiana* expression system with our LC-MS approach now enabled us to study substrate specificities of key enzymes in the PA network by simply adding stable isotope variants of arginine, ornithine, and lysine to plant extracts. These assays showed that transiently expressed tomato ADCs and ODC1 enzymes had the expected substrate specificities (Figure 6).

Overall, addition of a simple extraction procedure has opened the door for analyzing the key enzymes for PA synthesis, ADC and ODC, utilizing the ease of transient expression in *N. benthamiana*. This not only enables comparative studies of enzyme mutants or homologs, but also following different treatments (e.g., inhibitor, hormone, or abiotic/biotic stress treatments), all within the same *N. benthamiana* leaf background. LC-MS analysis of *N. benthamiana* leaves ectopically expressing enzymes of interest eliminates the need for time- and resource-intensive recombinant protein expression and purification, streamlining enzyme activity studies, making them faster and more accessible.

*N. benthamiana* is not only a commonly used system for *in planta* recombinant expression of proteins of interest but also serves as a host for the bacterial model pathogens *Pseudomonas* and *Xanthomonas*. The assays established here now enable the combination of transient expression of enzymes that alter PA homeostasis with infections by *Pseudomonas* or *Xanthomonas*, both of which have been shown to manipulate the PA pathway through injected toxins or effector proteins (Kim et al., 2013, Guo et al., 2023, Gerlin et al., 2021). Such assays, together with our MS-based platform for PA quantification, could provide a powerful tool to investigate how different PAs influence bacterial infection.

#### Broadening the scope of quantified metabolites provides insights into the wider PA network

We also aimed to broaden our portfolio of PA-related metabolites that can be quantified by our LC-MS approach, particularly focusing on anabolic pathway components, many of which play important roles in plant stress responses (Joshi and Fernie, 2017, Hildebrandt, 2018, Batista-Silva et al., 2019, Moormann et al., 2022). We compared 11 PA network members in RT and CT tomato leaves, finding a CT-dependent reduction of arginine and lysine levels and increased agmatine, putrescine, citrulline, and ornithine levels (Figures 2 and 8). The arginine pool was likely depleted due to increased *SlADC1/2* expression and ADC activity following CT (Supplementary Figures 1 and 9, Figures 2C and 8A), accumulating agmatine for putrescine synthesis, a metabolite known to mitigate cold stress (Cuevas et al., 2008, Chen et al., 2019, Upadhyay et al., 2020, Gonzalez-Hernandez et al., 2022). Interestingly, despite increased ADC activity, agmatine, and putrescine levels, we never observed CT-dependent changes in spermidine or spermine levels (Figures 3 and 8). Similar results have been previously reported (Alcázar et al., 2005, Alcazar et al., 2006, Wu et al., 2019, Gallas et al., 2024), indicating that spermidine and spermine levels are strictly regulated, possibly by PA-dependent translational regulation (Jimenez-Bremont et al., 2022). By displaying this large amount of analytical data as a pathway heatmap diagram, it is easy to spot metabolites whose homeostasis may be prioritized (i.e., spermidine and spermine) or diverted to other metabolic pathways during stress responses (e.g., lysine). Together, quantification of the wider PA network and visualization via a heatmap provide a comprehensive insight into PA pathway fluxes, illustrating genotype- or condition-specific pathway flux alterations and exposing strict regulatory points.

#### Insights into the roles of SlADCs on the PA network and plant development using CRISPR-Cas9-generated tomato mutants

We also used our optimized LC-MS-based approach to study the role of tomato ADC and ODC enzymes on ADC activity and PA metabolism following an acute CT using single *adc1* and *adc2* and double *adc1/adc2* tomato mutants (Wu et al., 2019) and a novel *odc1* mutant (Supplementary Figure 8). Importantly, we confirmed that SlADC1 and SlADC2 are the sole enzymes capable of synthesizing agmatine in tomato as no ADC activity was observed above background levels in the *adc1/adc2* double mutant (Figure 8A). Accordingly, compared to WT, this *adc1/adc2* mutant had higher arginine levels and lower agmatine and putrescine under both RT and CT conditions (Figure 8B). This demonstrates that in the absence of functional ADC enzymes, arginine builds up, and as agmatine cannot be produced, putrescine synthesis is reduced. As *adc1/adc2* plants have a functional ODC pathway but still lower putrescine levels compared to WT, this indicates that SlODCs cannot compensate for the loss of SlADCs. Surprisingly, no difference in spermidine or spermine levels were observed in the *adc1/adc2* mutant compared to WT in either RT or CT samples, despite reduced levels of their precursor putrescine (Figure 8B). This suggests that despite completely lacking the ADC-dependent pathway, spermidine and spermine levels are maintained, indicating a prioritization regulatory mechanism maintaining their levels at the expense of putrescine.

Alternatively, PAs may be synthesized in other tissues with higher ODC expression, such as the roots (Kwak and Lee, 2001, Acosta et al., 2005) and transported through the plant. Grafting the arial part of *adc1/adc2* plants to WT root stocks and analyzing ADC activity and PA levels in their leaves could give insight into long distance transport of such metabolites.

The *adc1* single mutant also had severely reduced ADC activity compared to *adc2* and WT in both RT and CT conditions (Figure 8A), indicating that ADC1 is the main contributor of ADC activity in tomato leaves. This is likely due to higher *SlADC1* expression in tomato leaves (Supplementary Figures 1 and 9), rather than SlADC1 being a more efficient enzyme, as we observed only slightly higher ADC activity levels when transiently overexpressing SlADC1 compared to SlADC2 in *N. benthamiana* (Figures 4, 5, and 6).

We also characterized the phenotypes of these single and double tomato mutants, as a double *adc1/adc2* mutant in *A. thaliana* cannot be generated (Urano et al., 2005). Compared to WT, *adc1/adc2* mutant plants were severely affected, being only two-thirds as tall as the WT at 47 DAG, with significantly fewer adult leaves and no flowers (Figure 7). In fact, we never observed any developing floral buds on *adc1/adc2* plants, even after months of growth. Interestingly, spermine has previously been found to be essential for flower development in *A. thaliana* (Zierer et al., 2016, Chen et al., 2019). Yet we observed no significant differences in spermidine and spermine levels in *adc1/adc2* leaves compared to WT (Figure 8). Therefore, at least in tomato, an as-yet unidentified ADC-dependent metabolite appears to be essential for flowering in tomato. Grafting experiments could clarify if a WT rootstock or stem could metabolically complement the *adc1/adc2* mutant and restore flower development. Similarly, exogenous agmatine supplementation of adult *adc1/adc2* plants or local T-DNA mediated transient expression *in planta* using a newly identified tomato-compatible *Rhizobium rhizogenes* strain (Lopez-Agudelo et al., 2025) may also facilitate flower development in this mutant. Overall, this work demonstrates that ADCs are essential for plant and flower development, even though the concentrations of key PAs, such as spermidine and spermine remain unchanged in the *adc1/adc2* mutant.

### Summary

In summary, we optimized our LC-MS-based method to quantify ADC and ODC enzyme activities and 11 PA-related metabolites in a single plant extract by a single MS run. This method has multiple advantages over other published methods: it is reasonably high throughput (manageable to extract and run up to 100 samples/day), less damaging to LC-MS systems (HFBA is not required in the LC-MS gradient), and does not require derivatization, reducing the overall cost and labor efforts of such an experiment. Furthermore, it is the only published method that measures both ADC and ODC enzyme activity and PA levels from the same sample. We believe our LC-MS-based method will be invaluable to researchers studying plant PA metabolism, including those investigating how plant pathogens manipulate this network via injected effectors or toxins (Wu et al., 2019, Gerlin et al., 2021). Given that ADCs are indispensable for plant reproduction and PAs play key roles in stress responses, our method could also be instrumental in advancing research areas across various fields of plant biology.

## AUTHOR CONTRIBUTIONS

ESR and EvR-L conducted experiments, EvR-L conducted metabolite profiling. DP generated heatmap plot and assisted with experiments. DW generated CRISPR-Cas9 tomato mutants and was involved in early method development. ESR, EvR-L, and TL wrote the manuscript.

## ACKNOWLEDGMENTS

We thank the ZMBP cultivation teams for plant transformation and care. Research in the Lahaye Laboratory on polyamines and bacterial effectors manipulating them is supported by DFG grants LA 1338/15-1, LA 1338/ 17-1, and TRR356 (project number: 491090170, subproject B03). Metabolite analytics in ZMBP central facilities unit was funded by the DFG (Project number 442641014). Research in the Wu laboratory is supported by the National Natural Science Foundation of China (32372485). ChatGPT (OpenAI, version 3.5) was used for mild text editing.

---

**Supplementary Figure 1:**
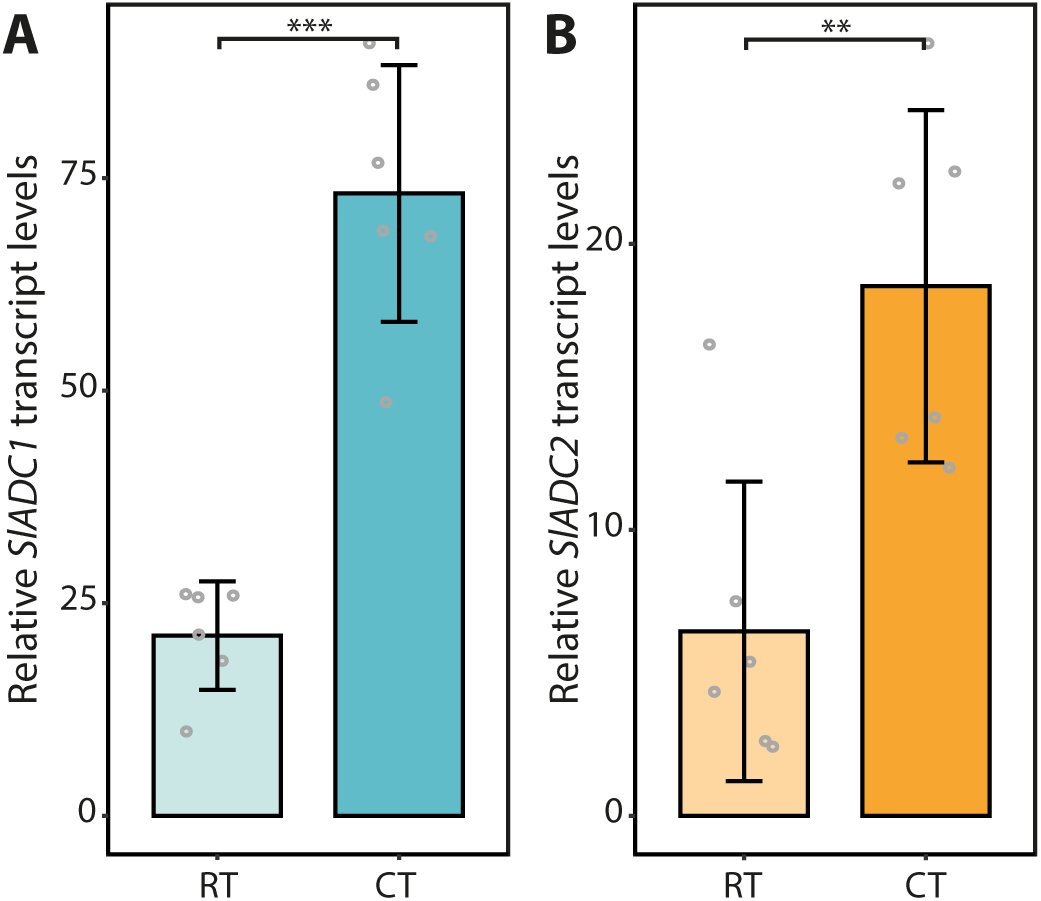
Cold treatment of WT adult tomato leaves induces SlADC1 and SlADC2 expression. RT-qPCR conducted on RNA extracted from 24-hour cold treated (CT) or room temperature (RT) control WT adult tomato leaves determining A) *SlADC1* and B) *SlADC2* transcript levels relative to the house keeping SlTIP41. Grey circles represent the mean of 3 technical replicates originating from one biological replicate (n =6). Statistical significance determined using a Student’s T Test (a = 0.05); * = P-value < 0.05, ** = P-value < 0.01, and *** = P-value < 0.001.

**Supplementary Figure 2:**
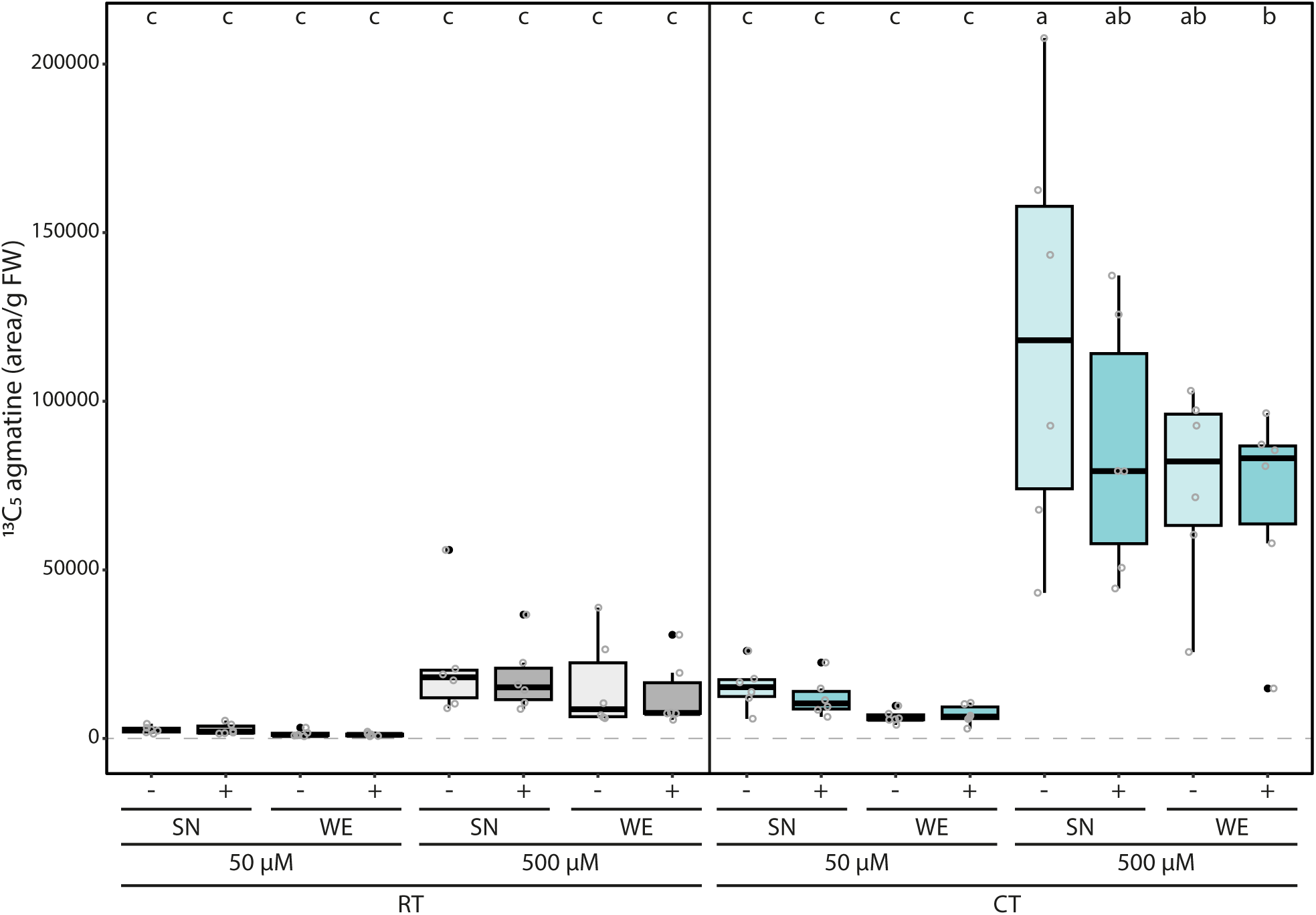
Inclusion of HFBA in LC-MS gradient does not affect ADC activity quantification. The same samples presented in Figure 2 that were run without HFBA in the LC-MS gradient solvent (-; light shades) were re-run with 0.1% HFBA included in the gradient (+; darker shades). ADC activity (13C5 agmatine in peak area/g fresh wight [FW] tissue, normalized to internal D5 tryptophan standard) in tomato leaf samples, pretreated either with room temperature (RT; grey) or cold treatment (CT; blue), with either 50 or 500 µM 13C6 arginine added to the SN or WE. Statistical significance determined using ANOVA followed by Tukey’s post hoc HSD test; different letters above each box indicate statistically significantly groups (a = 0.05). Each biological replicate represented by a grey circle; n = 6.

**Supplementary Figure 3:**
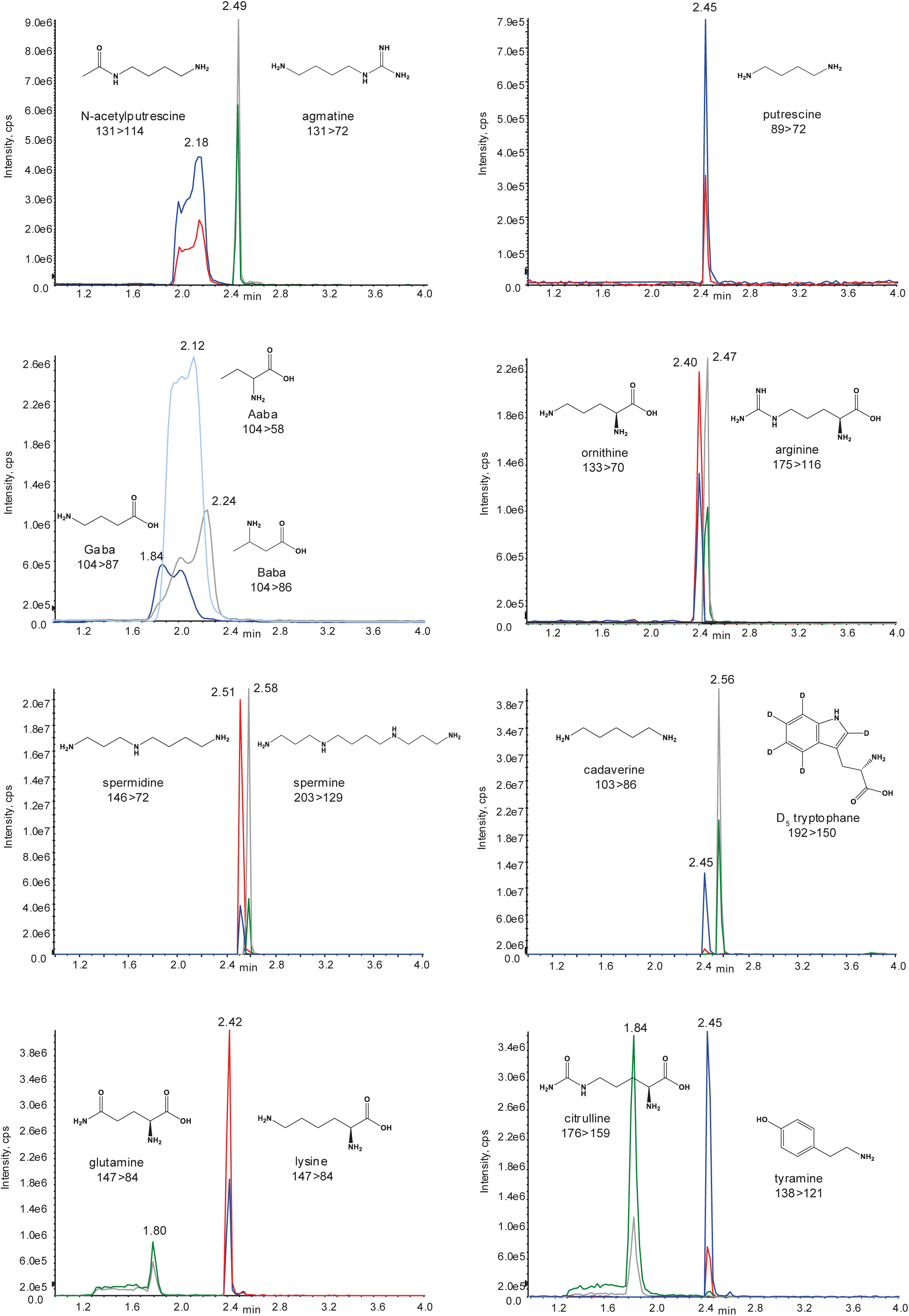
Extracted ion chromatograms (XICs) of members of the PA network. Zoomed in XICs (1-4 minutes of the 5 min gradient; counts per second [cps]), retention times (RT; min), and chemical structures of N-acetylputrescine, agmatine, putrescine, Gaba, Aaba, Baba, ornithine, arginine, spermidine, spermine, cadaverine, D5 tryptophane, glutamine, lysine, citrulline, and tyramine. All standards were run together in a concentration of 100 nM, except the aminobutyric acid variants (1000 nM) and glutamine (200 nM), diluted in water with 0.1% formic acid and 0.1% HFBA. In most cases two MRM transitions are shown with the quantifier ion stated in the chromatogram (red or grey XIC). Additional information (LOQ, linearity range, qualifier ion transitions) can be found in Supplementary Table 1.

**Supplementary Figure 4:**
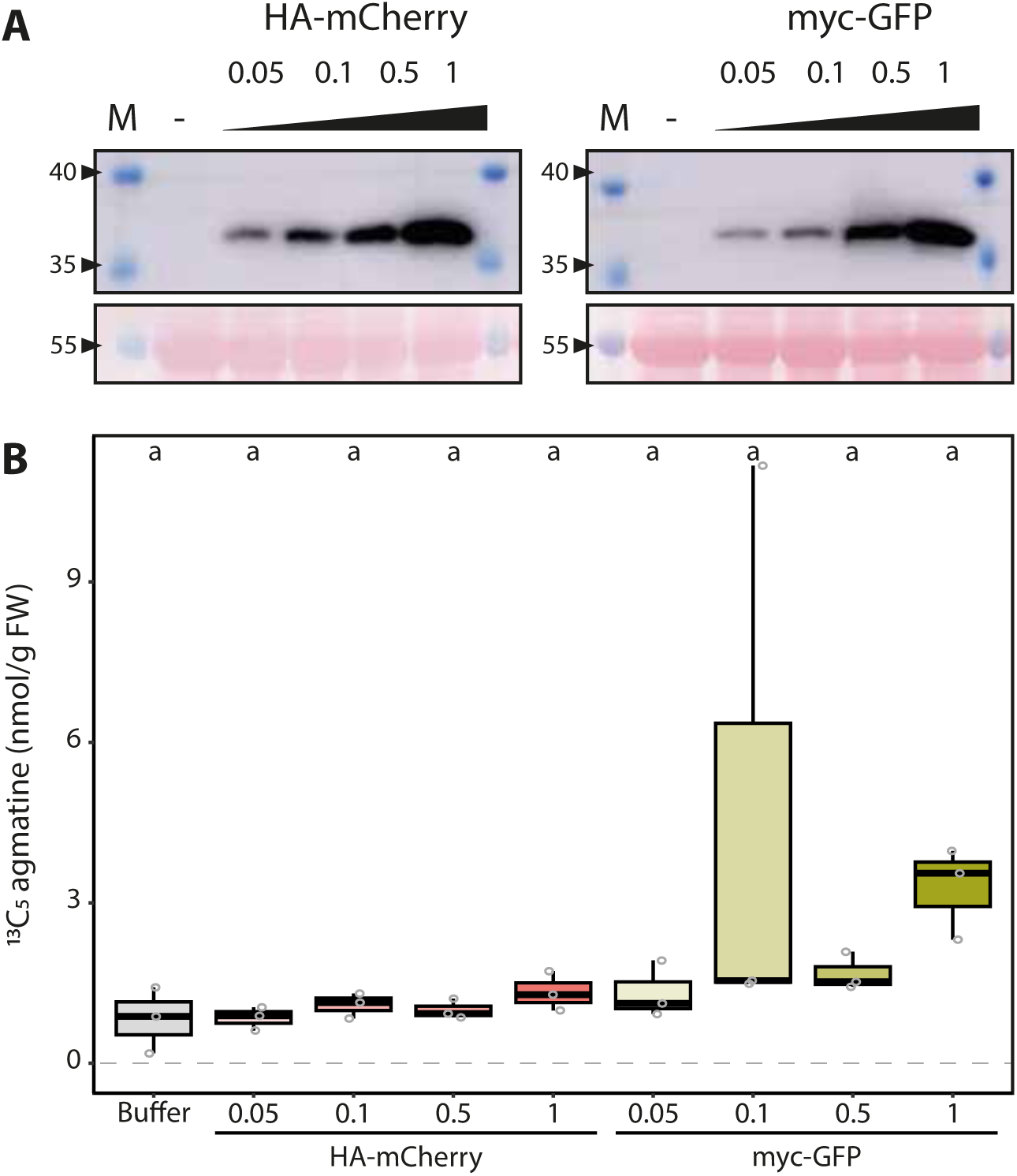
Increasing amounts of fluorophore negative controls do not linearly correlate with rise in ADC activity. **A)** Protein abundance of fluorophore controls increases with increasing bacterial load. Immunoblot analysis of *N. benthamiana* leaves agroinfiltrated with 35Sp:HA-mCherry:35St (left) and 35Sp:myc-GFP:35St (right) at increasing OD600 (0.05, 0.1, 0.5, and 1) of samples taken 2 days post infiltration (dpi). Samples of leaves infiltrated with only infiltration buffer were included as a negative control (-). Upper images (cropped) show HA-mCherry detection (left) and myc-GFP (right); expected size for both proteins is 29 kDa. Ponceau stained blots show equal protein loading (bottom). **B)** ADC activity does not correlate with increasing fluorophore protein abundance. ADC activity (13C5 agmatine in nmol/g FW, normalized to D5 tryptophan) from the same *N. benthamiana* sample agroinfiltrated with HA-mCherry (red) or myc-GFP (green), or infiltration buffer only (grey) as shown in **A)**. Increasing color intensity represents increasing bacterial loads. Statistical significance determined using ANOVA followed by Tukey’s post hoc HSD test; different letters above each box indicate statistically significantly different groups (a = 0.05 level). Each biological replicate represented by a grey circle; n = 3.

**Supplementary Figure 5:**
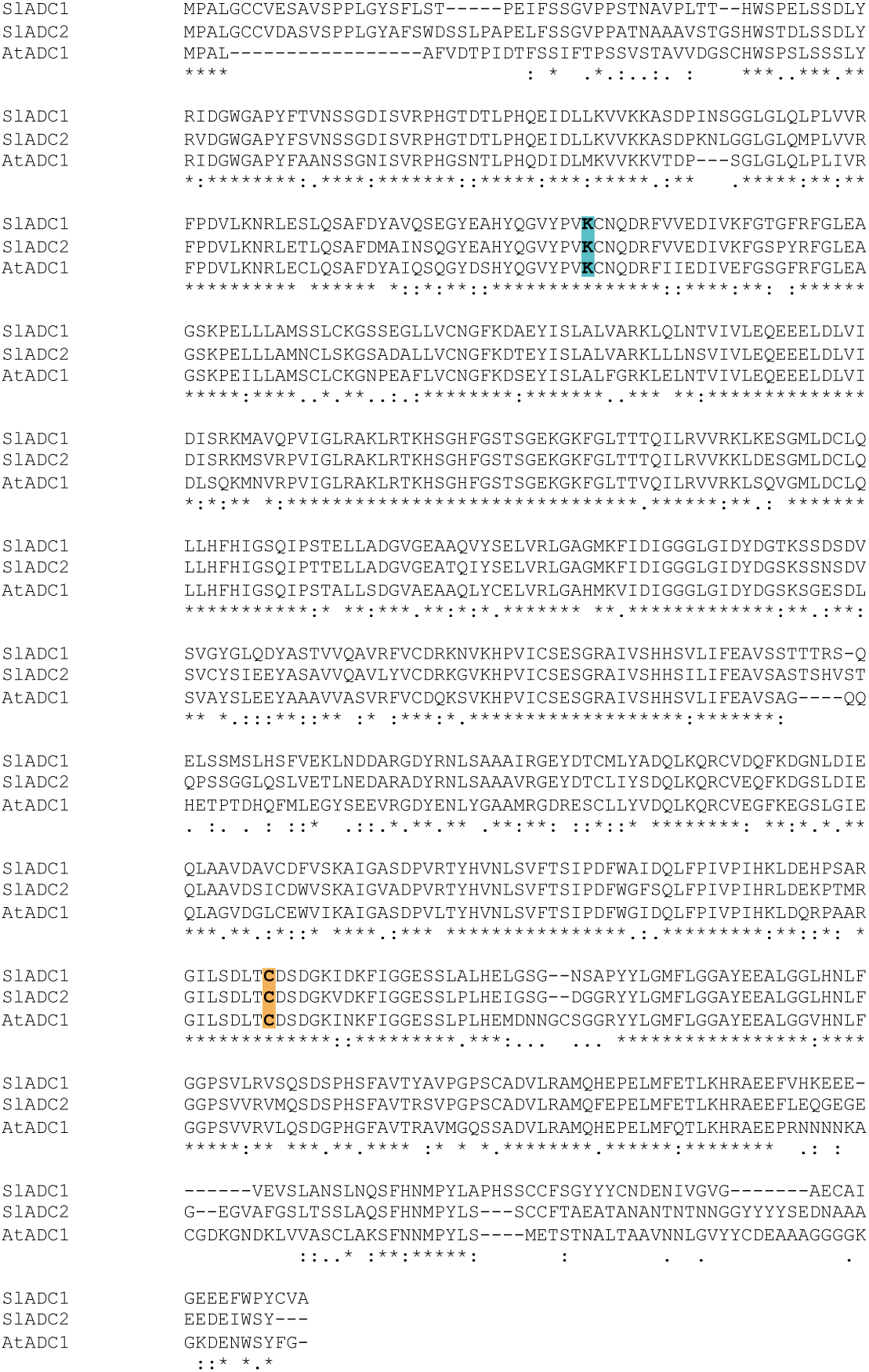
Alignment of tomato ADC1 and ADC2 and *A. thaliana* ADC1 peptide sequences, highlighting essential catalytic residues. Peptide sequences of tomato ADC1 (SlADC1), tomato ADC2 (SlADC2), and *A. thaliana* ADC1 (AtADC1) were aligned using CLUSTALW online tool (https://www.genome.jp/tools-bin/clustalw). Homology between sequences is indicated below alignment, where an asterisk (*) indicates identical residues for all sequences, while dots indicate only partial conservation (. or :). Conserved lysine (K; blue) binds ADC co-factor, pyridoxal 5-phosphate. Conserved cysteine (C; orange) binds ADC substrate.

**Supplementary Figure 6:**
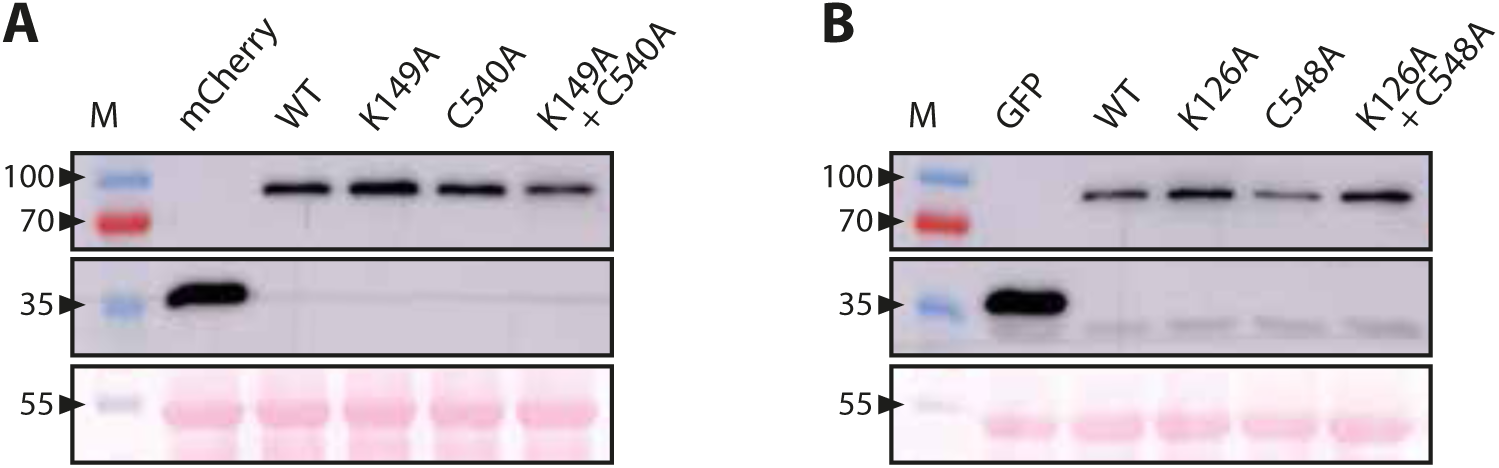
All WT and mutant tomato ADC proteins can be transiently expressed in *N. benthamiana* leaves. **A)** Immunodetection of transiently expressed SlADC1 WT and mutant variants in N. benthamiana leaves at 2 dpi. HA epitope tagged tomato ADC1 WT, K149A, C540A, and the double mutant K149A + C540A T-DNA constructs were agroinfiltrated into WT *N. benthamiana* leaves alongside the negative control HA-mCherry. Western blot analysis enabled detection of proteins using an anti-HA-horseradish peroxidase (HRP)-conjugated antibody. **B)** Immunodetection of transiently expressed SlADC2 WT and mutant variants in *N. benthamiana* leaves at 2 dpi. Epitope-tagged (myc) tomato ADC2 WT, K126A, C548A, and the double mutant K126A + C548A T-DNA constructs were agroinfiltrated into WT N. benthamiana leaves alongside the negative control, myc-GFP. Detection of proteins using an anti myc primary antibody and conjugated anti mouse HRP secondary antibody. Expected protein sizes are 78 kDa, 80 kDa, and 29 kDa for HA-vSlADC1 variants, myc-SlADC1 variants, and the tagged fluorophores, respectively. Ponceau stained blots show equal protein loading (below each western blot).

**Supplementary Figure 7:**
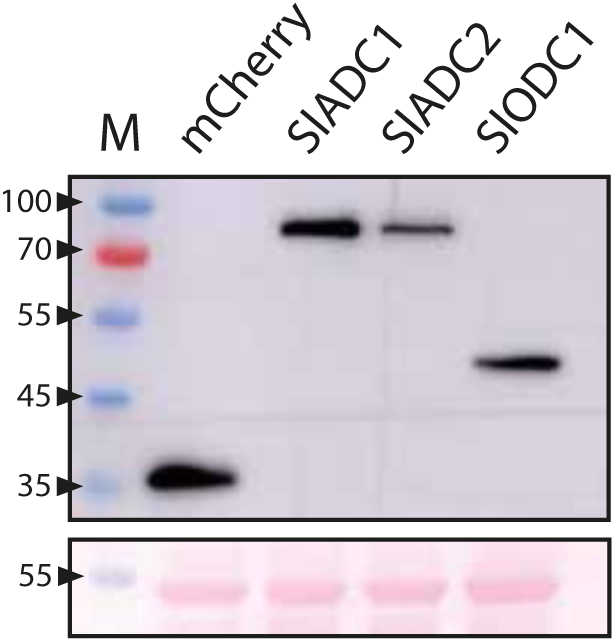
Tomato ADC1, ADC2, and ODC1 proteins are expressed in *N. benthamiana* leaves. Immunodetection of HA epitope-tagged tomato ADC1, ADC2, and ODC1 enzymes and the negative mCherry control transiently expressed in *N. benthamiana* leaves. After 2dpi, samples were harvested for western blot analysis using an anti HA horseradish peroxidase (HRP) conjugated antibody to detect proteins. Expected protein sizes are 78 kDa, 80 kDa, 48 kDa, and 29 kDa for HA-SlADC1, HA-SlADC2, HA-SlODC1, and HA-mCherry, respectively. Ponceau stained blots show equal protein loading (below).

**Supplementary Figure 8:**
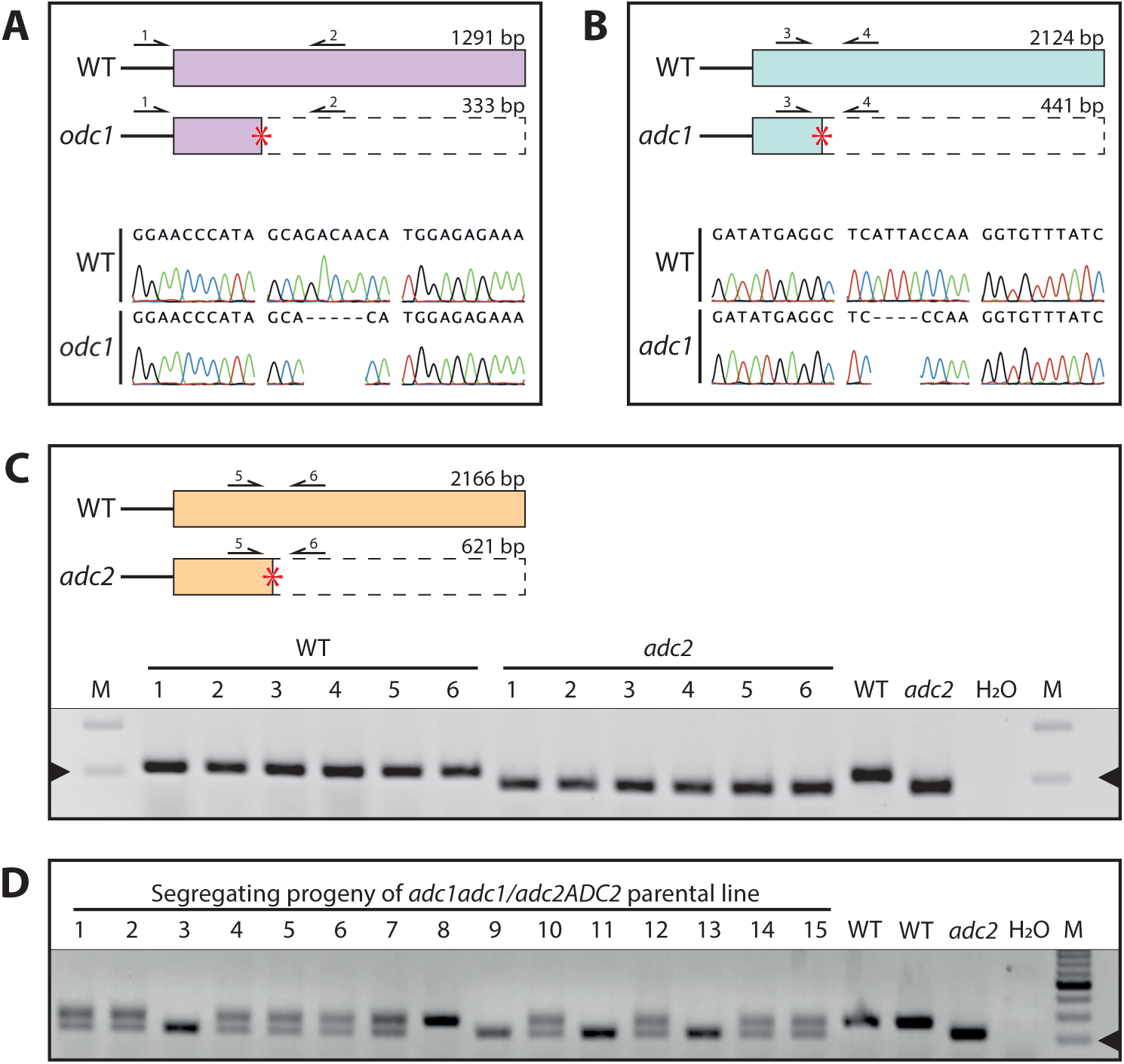
CRISPR Cas9 mutagenesis of tomato ODC1, SlADC1, and SlADC2 genes to generate knock out mutants. **A)** CRISPR-Cas9 mutagenesis of tomato *odc1* (Sl04g082030.1) resulted in 5 base pair (bp) deletion, causing a frameshift and premature stop codon. Schematic of WT and *odc1* mutant coding sequence (CDS) structure showing premature stop codon (red asterisks) after 333 bp (top). Genotyping was conducted by PCR amplification from extracted gDNA using primers 1 and 2 (black arrows) and sequencing PCR products; trace data from WT and *odc1* mutant shows 5 bp deletion (bottom). **B)** Mutagenesis of *SlADC1* (Sl10g054440.2) resulted in a 4 bp deletion, causing a frameshift and premature stop codon after 441 bp (top). Genotyping was conducted in the same manner with primers 3 and 4; trace data shows 4 bp deletion (bottom). **C)** CRISPR-Cas9 mutagenesis of *SlADC2* (Sl01g110440.4) produced a 52 bp deletion. Schematic of WT and *adc2* mutant CDS structure shows premature stop codon after 621 bp (top). Genotyping was conducted via PCR amplification using primers 5 and 6; amplification of WT produces 272 bp PCR product, while *adc2* produces 220 bp. Previously verified WT and *adc2* genotypes were used as controls, alongside water (H2O) as a negative PCR control. Black arrow indicates 250 bp band in marker lane. **D)** Generation of *adc1/adc2* double mutant requires genotyping segregating progeny of parental adc1adc1/adc2ADC2 line. Homozygous, Cas9 transgene-free adc1 and adc2 mutants (shown in B and C) were crossed and an *adc1adc1/adc2ADC2* line in the F2 generation was selected. To obtain sufficient *adc1/adc2* plants for phenotyping (i.e., n = 20; Figure 7), 200 progeny plants of this *adc1adc1/adc2ADC2* line were genotyped via PCR amplification of *SlADC2* CDS using primers 5 and 6; typically, only ∼10% of progeny pants are adc1/adc2 when using recently harvested seed. Using previously verified WT and *adc2* genotypes as controls, progeny plants homozygous for the *adc2* deletion (i.e., band size 220 bp, verified in two replicate PCR experiments) were selected as *adc1/adc2* plants; for example, plants 3, 9, 11, and 13 here. Black arrow indicates 200 bp band in marker lane.

**Supplementary Figure 9:**
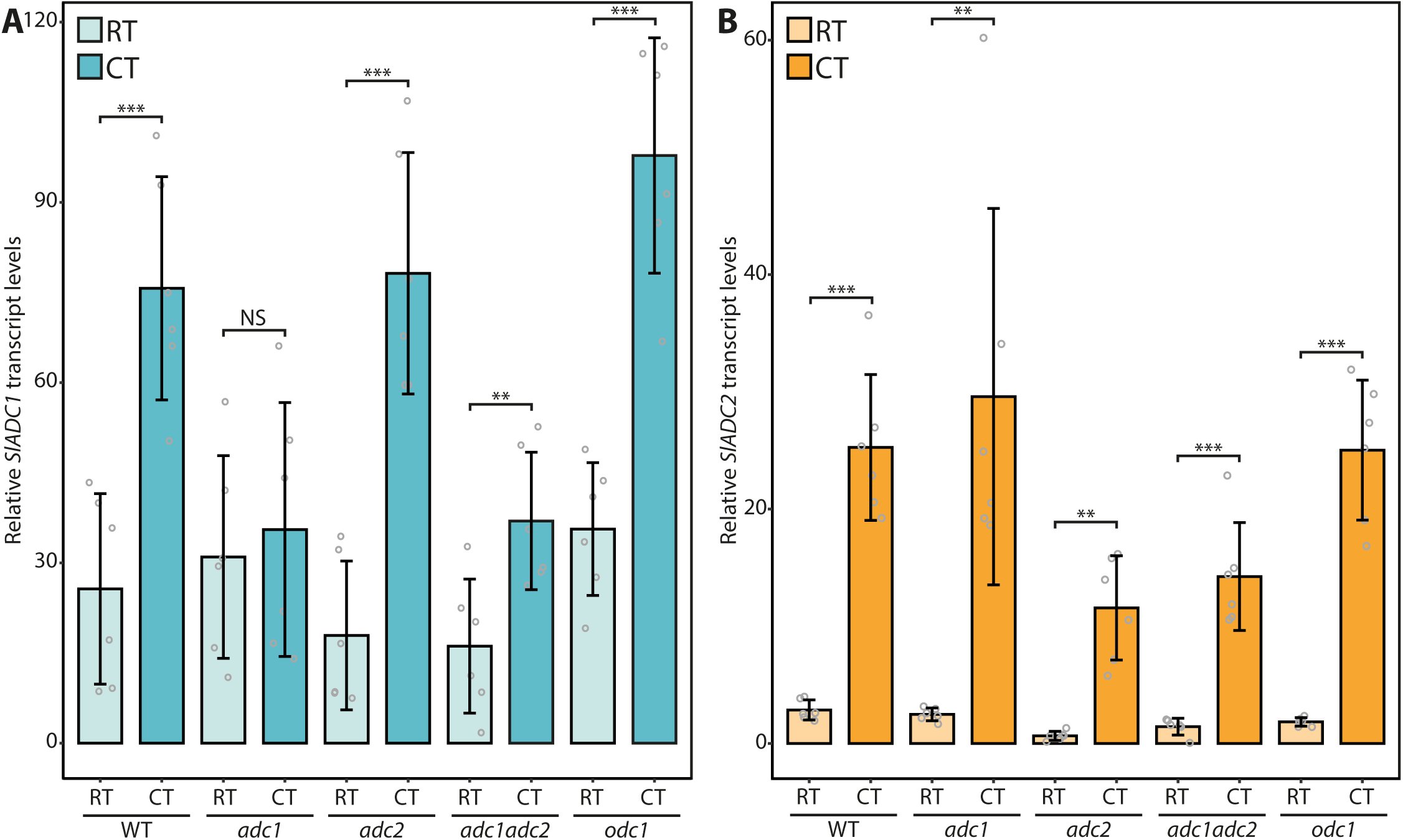
Cold treatment of WT, *adc1*, *adc2*, *adc1/adc2*, and *odc1* adult tomato leaves induces SlADC1 and *SlADC2* expression. RNA extracted from WT *adc1*, *adc2*, *adc1/adc2*, and *odc1* tomato leaves that were cold treated (CT) for 24 hours or the room temperature (RT) controls was used for RT-qPCR to determine **A)** *SlADC1* and **B)** *SlADC2* transcript levels relative to the house keeping SlTIP41 gene expression. Grey circles represent the mean of 3 technical replicates originating from the 6 biological replicates. Statistical significance determined using a Student’s T Test to compare expression between each condition from each genotype (a = 0.05); * = P-value < 0.05, ** = P-value < 0.01, and *** = P-value < 0.001.

**Supplementary Figure 10.**
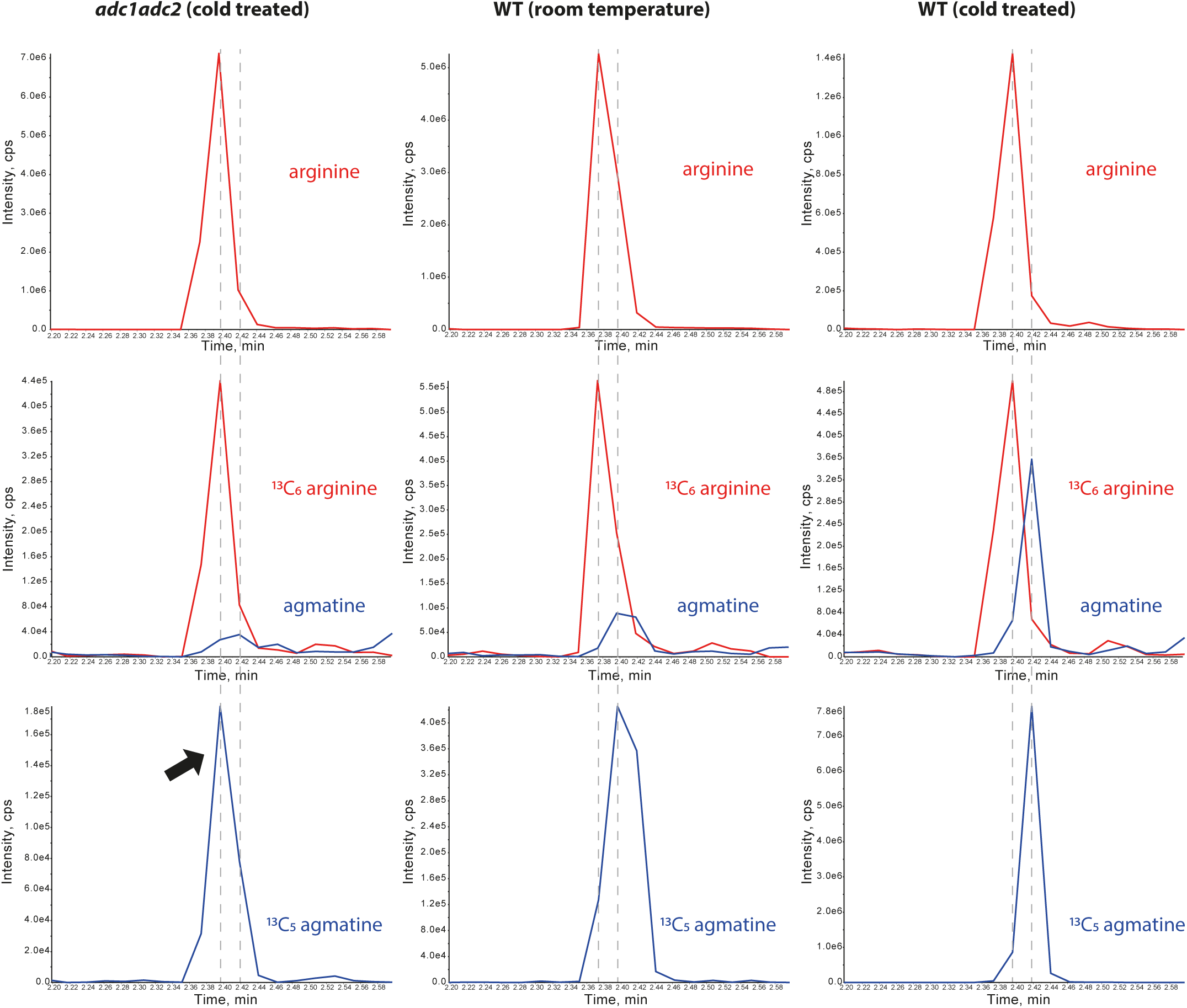
Extracted ion chromatograms (XICs) of 13C5 agmatine in *adc1/adc2* following cold treatment confirm lack of ADC activity. XICs of cold treated (CT) *adc1/adc2* double mutant extracts (left column) showed only traces of 13C5 agmatine (bottom row; black arrow), but this peak was congruent to arginine (red; top row) and 13C6 arginine peaks (red; middle row; measured on the second isotope). Room temperature (RT) and CT treated WT plant extracts (middle and right column) show enzymatically produced 13C5 agmatine (blue), which has a peak lining up to agmatine peaks (blue; middle panel). Dotted grey lines indicate either arginine or agmatine peaks in each plant extract.

## Supplementary tables

**Supplementary Table 1:** Transitions (Q1/Q3), declustering potential (DP), collision energy (CE), retention times (RT), limit of detection (LOD), limit of quantitation (LOQ) and the linear range for the recorded analytes

|  |  |  |  |  |  |  |  |  |
| --- | --- | --- | --- | --- | --- | --- | --- | --- |
| Q1 mass (Da) | Q3 mass (Da) | DP (volts) | CE (volts) | Compound | RT min | LOD nM | LOQ nM | Linear range nM |
| 175.1 | 158.0 | 50 | 20 | arginine 1 | 2.47 |  |  |  |
| **175.1** | **116.1** | 50 | 20 | arginine 2 |  | 0.1 | 1 | 1-10 |
| 181.1 | 164.1 | 50 | 20 | <sup>13</sup>C<sub>6</sub> arginine 1 | 2.47 |  |  |  |
| **181.1** | **121.1** | 50 | 20 | <sup>13</sup>C<sub>6</sub> arginine 2 |  |  |  |  |
| **182.1** | **122.1** | 50 | 20 | <sup>13</sup>C<sub>6</sub> arginine 2 second isotopes * | 2.47 |  |  |  |
| 131.1 | 114.1 | 22 | 15 | agmatine 1 | 2.49 | 0.5 | 5 | 5-50 |
| **131.1** | **72.1** | 22 | 20 | agmatine 2 |  |  |  |  |
| 136.1 | 119.1 | 22 | 15 | <sup>13</sup>C<sub>5</sub> agmatine 1 | 2.49 |  |  |  |
| **136.1** | **76.1** | 22 | 20 | <sup>13</sup>C<sub>5</sub> agmatine 2 |  |  |  |  |
| 133.0 | 116.0 | 50 | 10 | ornithine 1 | 2.40 |  |  |  |
| **133.0** | **70.0** | 50 | 20 | ornithine 2 |  | 0.1 | 1 | 1-100 |
| 140.1 | 122.1 | 50 | 10 | <sup>13</sup>C<sub>5</sub><sup>15</sup>N<sub>2</sub> ornithine 1 | 2.40 |  |  |  |
| **140.1** | **75.0** | 50 | 20 | <sup>13</sup>C<sub>5</sub><sup>15</sup>N<sub>2</sub> ornithine 2 |  |  |  |  |
| **141.1** | **76.0** | 50 | 20 | <sup>13</sup>C<sub>5</sub><sup>15</sup>N<sub>2</sub> ornithine 2 second isotopes * | 2.40 |  |  |  |
| **89.1** | **72** | 22 | 14 | putrescine 1 | 2.45 | 1 | 1 | 5-100 |
| 89.1 | 72 | 22 | 25 | putrescine 2 |  |  |  |  |
| **93.1** | **76.1** | 22 | 14 | <sup>13</sup>C<sub>4</sub> putrescine | 2.45 |  |  |  |
| **95.1** | **77.1** | 22 | 14 | <sup>13</sup>C<sub>4</sub><sup>15</sup>N<sub>2</sub> putrescine | 2.45 |  |  |  |
| **131.1** | **114.1** | 22 | 15 | N-acetylputrescine 1 | 2.18 | 0.5 | 1 | 1-100 |
| 131.1 | 72.0 | 22 | 20 | N-acetylputrescine 2 |  |  |  |  |
| 146.15 | 129 | 20 | 20 | spermidine 1 | 2.51 |  |  |  |
| **146.15** | **72** | 20 | 20 | spermidine 2 |  | 0.1 | 1 | 1-50 |
| 203.2 | 112 | 48 | 15 | spermine 1 | 2.58 |  |  |  |
| **203.2** | **129.1** | 48 | 24 | spermine 2 |  | 0.1 | 10 | 10-250 |
| **176.1** | **159.1** | 22 | 15 | citrulline 1 | 1.84 | 0.1 | 0.5 | 0.5-250 |
| 176.1 | 113.1 | 22 | 25 | citrulline 2 |  |  |  |  |
| **179.1** | **162.1** | 22 | 15 | <sup>13</sup>C<sub>1</sub>D<sub>2</sub> citrulline 1 | 1.84 | 0.1 | 0.5 | 0.5-250 |
| 179.1 | 116.2 | 22 | 20 | <sup>13</sup>C<sub>1</sub>D<sub>2</sub> citrulline 2 |  |  |  |  |
| 147.1 | 130.0 | 50 | 15 | lysine 1 | 2.42 |  |  |  |
| **147.1** | **84.0** | 50 | 25 | lysine 2 |  | 1 | 1 | 1-50 |
| **156.1** | **137.1** | 50 | 15 | <sup>13</sup>C<sub>6</sub><sup>15</sup>N<sub>2</sub> lysine 2 second isotopes * | 2.42 |  |  |  |
| **103.1** | **86.0** | 40 | 25 | cadaverine 1 | 2.45 | 0.1 | 0.1 | 0.1-10 |
| 103.1 | 69.1 | 40 | 25 | cadaverine 2 |  |  |  |  |
| **110.1** | **92.0** | 40 | 15 | <sup>13</sup>C<sub>5</sub><sup>15</sup>N<sub>2</sub> cadaverine 1 | 2.45 |  |  |  |
| 110.1 | 74.1 | 40 | 25 | <sup>13</sup>C<sub>5</sub><sup>15</sup>N<sub>2</sub> cadaverine 2 |  |  |  |  |
| **147.1** | **130.0** | 40 | 10 | glutamine 1 | 1.80 | 10 | 10 | 10-500 |
| 147.1 | 84.0 | 40 | 30 | glutamine 2 |  |  |  |  |
| **104.1** | **87.0** | 90 | 17 | γ-aminobutyric acid (GABA) 1 | 1.84 | 10 | 10 | 10-500 |
| **104.1** | **86.0** | 90 | 13 | β-aminobutyric acid (BABA) 1 | 2.24 |  |  |  |
| **104.1** | **58.0** | 90 | 19 | α-aminobutyric acid (AABA) 1 | 2.12 |  |  |  |
| **138.1** | **121.1** | 40 | 15 | tyramine 1 | 2.45 | 0.1 | 1 | 1.50 |
| 138.1 | 103.1 | 40 | 30 | tyramine 2 |  |  |  |  |
| 210.1 | 192 | 60 | 22 | D<sub>5</sub> tryptophane 1 | 2.56 |  |  |  |
| **192.1** | **150.1** | 100 | 22 | D<sub>5</sub> tryptophane 2 |  | 0.1 | 0.1 | 0.1-50 |

Multiple reaction monitoring (MRM) transitions in bold letters represent the quantifier ions. Limits of detection (LOD) and limits of quantitation (LOQ) were determined based on visual evaluation, signal-to-noise (S/N) determination, and on the standard deviation of the response and calibration curve slope. S/N was calculated using the relative noise and autpeak integration algorithm of the Sciex OS software. For evaluating the linear range of the calibration curves, a mastermix containing the listed compounds above (with the exception of <sup>13</sup>C<sub>5</sub> agmatine, <sup>13</sup>C<sub>4</sub> putrescine, <sup>13</sup>C<sub>5</sub><sup>15</sup>N<sub>2</sub> putrescine, and <sup>13</sup>C<sub>5</sub><sup>15</sup>N<sub>2</sub> cadaverine) was diluted accordingly in the range of the LOD and LOQ in water with 0.1% HFBA and 0.1% FA. \* The second isotope transitions of <sup>13</sup>C<sub>6</sub> arginine, <sup>13</sup>C<sub>5</sub><sup>15</sup>N<sub>2</sub> ornithine, and <sup>13</sup>C<sub>6</sub><sup>15</sup>N<sub>2</sub> lysine were only monitored to check for substrate presence in the enzyme assays. The lower response pairs were chosen to prevent detector blinding of the MS system while using substrate levels in the μM range.

**Supplementary Table 2:** Chromatographic separation and LC-MS settings

| Supplementary Table 2: Chromatographic separation and LC-MS settings |  |  |
| --- | --- | --- |
| time: main separation gradient | % solvent A<br>(water, 0.1% aq. FA, + / - 0.05% HFBA) | % solvent B<br>(acetonitrile, 0.1% FA, + / - 0.05% HFBA) |
| 0 | 85 | 15 |
| 0.2 | 85 | 15 |
| 1.5 | 30 | 70 |
| 2.5 | 5 | 95 |
| 3 | 5 | 95 |
| 4 | 85 | 15 |
| 5 | 85 | 85 |
| time: trap gradient - HFBA | % solvent A<br>(water, 0.1% aq. FA) | % solvent B<br>(acetonitrile, 0.1% FA) |
| 0 | 98 | 2 |
| 1.8 change to separation gradient | 98 | 2 |
| 4 | 98 | 2 |
| time: trap gradient + HFBA | % solvent A<br>(water, 0.1% aq. (FA), 0.05% HFBA) | % solvent B<br>(acetonitrile, 0.1% FA, 0.05% HFBA) |
| 0 | 98 | 2 |
| 2.5 change to separation gradient | 98 | 2 |
| 2.7 | 98 | 2 |

Flow rate 16 µl/min for the separation gradient, 25 µl/min for the trap gradients. Column temperature 50°C. MS settings: Optiflow Turbo V ion source with SteadySpray T micro electrode (10–50 µl/min); ion spray voltage: +4800 V; nebuliser, heater gas = nitrogen, 25 and 45 psi; curtain gas, nitrogen, 30 psi; collision gas, nitrogen, medium; source temperature, 200 °C; entrance potential, ±10 V; collision cell exit potential, ±10 V; dwell time 5 ms.

**Supplementary Table 3:**
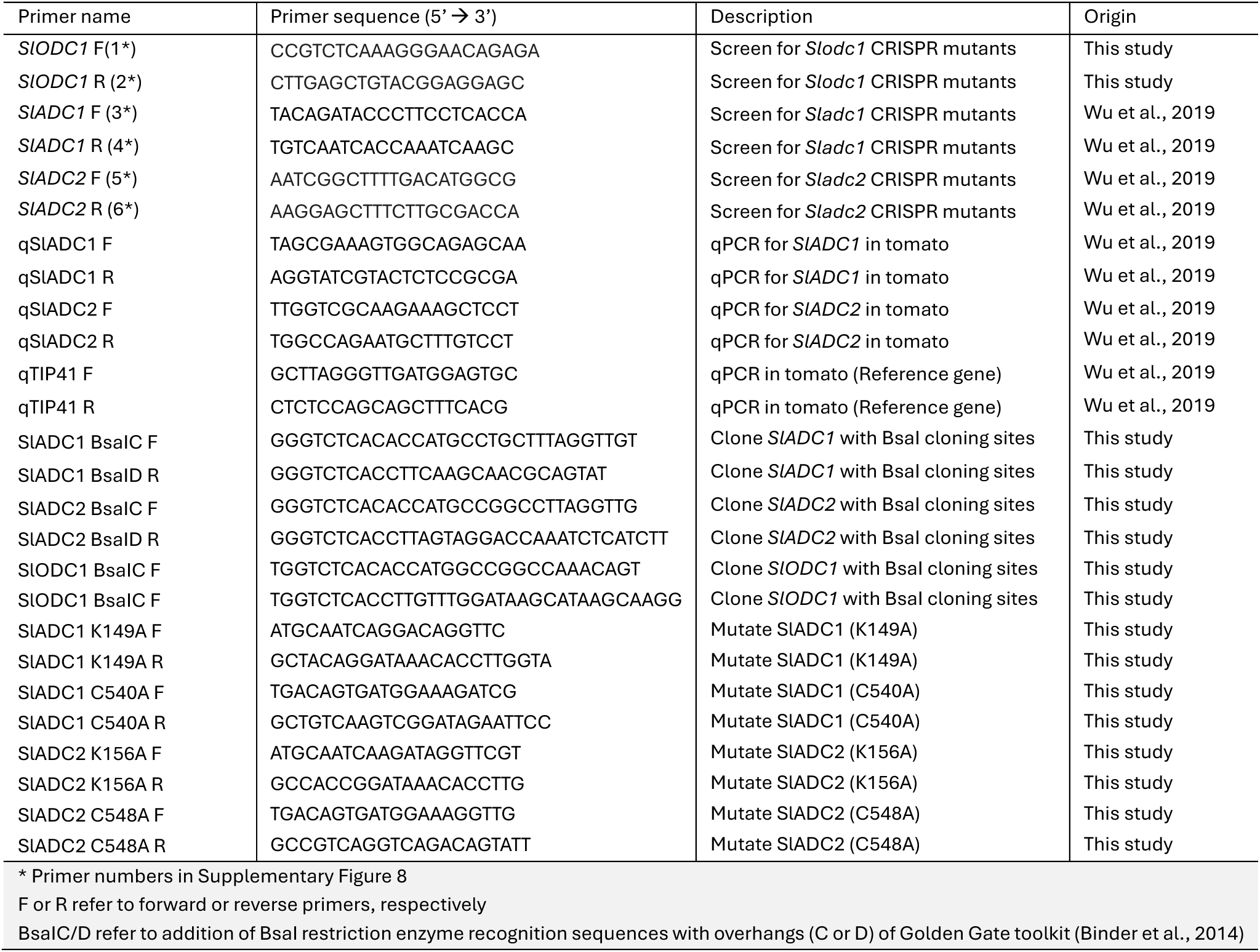
Primers used in this work

## REFERENCES

1. Acosta, C., Pérez-Amador, M. A., Carbonell, J. & Granell, A. 2005. The two ways to produce putrescine in tomato are cell-specific during normal development. Plant Science, 168, 1053–1057.

2. Alcazar, R., Altabella, T., Marco, F., Bortolotti, C., Reymond, M., Koncz, C., Carrasco, P. & Tiburcio, A. F. 2010. Polyamines: molecules with regulatory functions in plant abiotic stress tolerance. Planta, 231, 1237–49.

3. Alcázar, R., García-Martínez, J. L., Cuevas, J. C., Tiburcio, A. F. & Altabella, T. 2005. Overexpression of ADC2 in Arabidopsis induces dwarfism and late-flowering through GA deficiency. Plant Journal, 43, 425–436.

4. Alcazar, R., Marco, F., Cuevas, J. C., Patron, M., Ferrando, A., Carrasco, P., Tiburcio, A. F. & Altabella, T. 2006. Involvement of polyamines in plant response to abiotic stress. Biotechnol Lett, 28, 1867–76.

5. Bassard, J. E., Ullmann, P., Bernier, F. & Werck-Reichhart, D. 2010. Phenolamides: bridging polyamines to the phenolic metabolism. Phytochemistry, 71, 1808–24.

6. Batista-Silva, W., Heinemann, B., Rugen, N., Nunes-Nesi, A., Araujo, W. L., Braun, H. P. & Hildebrandt, T. M. 2019. The role of amino acid metabolism during abiotic stress release. Plant Cell Environ, 42, 1630–1644.

7. Binder, A., Lambert, J., Morbitzer, R., Popp, C., Ott, T., Lahaye, T. & Parniske, M. 2014. A Modular Plasmid Assembly Kit for Multigene Expression, Gene Silencing and Silencing Rescue in Plants. Plos One, 9.

8. Blázquez, M. A. 2024. Polyamines: Their Role in Plant Development and Stress. Annual Review of Plant Biology, 75, 95–117.

9. Bouchereau, A., Guénot, P. & Lather, F. 2000. Analysis of amines in plant materials. Journal of Chromatography B-Analytical Technologies in the Biomedical and Life Sciences, 747, 49–67.

10. Bunsupa, S., Katayama, K., Ikeura, E., Oikawa, A., Toyooka, K., Saito, K. & Yamazaki, M. 2012. Lysine decarboxylase catalyzes the first step of quinolizidine alkaloid biosynthesis and coevolved with alkaloid production in leguminosae. Plant Cell, 24, 1202–16.

11. Chen, D. D., Shao, Q. S., Yin, L. H., Younis, A. & Zheng, B. S. 2019. Polyamine Function in Plants: Metabolism, Regulation on Development, and Roles in Abiotic Stress Responses. Frontiers in Plant Science, 9.

12. Cuevas, J. C., Lopez-Cobollo, R., Alcazar, R., Zarza, X., Koncz, C., Altabella, T., Salinas, J., Tiburcio, A. F. & Ferrando, A. 2008. Putrescine is involved in Arabidopsis freezing tolerance and cold acclimation by regulating abscisic acid levels in response to low temperature. Plant Physiology, 148, 1094–1105.

13. Flores, H. E. & Galston, A. W. 1982. Analysis of Polyamines in Higher-Plants by High-Performance Liquid-Chromatography. Plant Physiology, 69, 701–706.

14. Gallas, N., Li, X., von Roepenack-Lahaye, E., Schandry, N., Jiang, Y., Wu, D. & Lahaye, T. 2024. An ancient cis-element targeted by Ralstonia solanacearum TALE-like eVectors facilitates the development of a promoter trap that could confer broad-spectrum wilt resistance. Plant Biotechnol J, 22, 602–616.

15. Gerlin, L., Baroukh, C. & Genin, S. 2021. Polyamines: double agents in disease and plant immunity. Trends Plant Sci, 26, 1061–1071.

16. Golubova, D., Tansley, C., Su, H. & Patron, N. J. 2024. Engineering as a platform for natural product biosynthesis. Current Opinion in Plant Biology, 81.

17. Gonzalez, M. E., Jasso-Robles, F. I., Flores-Hernandez, E., Rodriguez-Kessler, M. & Pieckenstain, F. L. 2021. Current status and perspectives on the role of polyamines in plant immunity. Annals of Applied Biology, 178, 244–255.

18. Gonzalez-Hernandez, A. I., Scalschi, L., Vicedo, B., Marcos-Barbero, E. L., Morcuende, R. & Camanes, G. 2022. Putrescine: A Key Metabolite Involved in Plant Development, Tolerance and Resistance Responses to Stress. Int J Mol Sci, 23.

19. Guo, Q., Chen, X. & Li, B. 2023. Purification and characterization of tomato arginine decarboxylase and its inhibition by the bacterial small molecule phevamine A. Protein Expr Purif, 210, 106326.

20. Hakkinen, M. R., Keinanen, T. A., Vepsalainen, J., Khomutov, A. R., Alhonen, L., Janne, J. & Auriola, S. 2007. Analysis of underivatized polyamines by reversed phase liquid chromatography with electrospray tandem mass spectrometry. J Pharm Biomed Anal, 45, 625–34.

21. Hanfrey, C., Sommer, S., Mayer, M. J., Burtin, D. & Michael, A. J. 2001. Arabidopsis polyamine biosynthesis: absence of ornithine decarboxylase and the mechanism of arginine decarboxylase activity. Plant J, 27, 551–60.

22. Heinemann, B. & Hildebrandt, T. M. 2021. The role of amino acid metabolism in signaling and metabolic adaptation to stress-induced energy deficiency in plants. J Exp Bot, 72, 4634–4645.

23. Hildebrandt, T. M. 2018. Synthesis versus degradation: directions of amino acid metabolism during Arabidopsis abiotic stress response. Plant Mol Biol, 98, 121–135.

24. Jacobs, T. B., Lafayette, P. R., Schmitz, R. J. & Parrott, W. A. 2015. Targeted genome modifications in soybean with CRISPR/Cas9. BMC Biotechnol, 15, 16.

25. Jancewicz, A. L., Gibbs, N. M. & Masson, P. H. 2016. Cadaverine’s Functional Role in Plant Development and Environmental Response. Front Plant Sci, 7, 870.

26. Jessome, L. L. & Volmer, D. A. 2006. Ion suppression: A major concern in mass spectrometry. Lc Gc North America, 83–89.

27. Jimenez-Bremont, J. F., Chavez-Martinez, A. I., Ortega-Amaro, M. A., Guerrero-Gonzalez, M. L., Jasso-Robles, F. I., Maruri-Lopez, I., Liu, J. H., Gill, S. S. & Rodriguez-Kessler, M. 2022. Translational and post-translational regulation of polyamine metabolic enzymes in plants. J Biotechnol, 344, 1–10.

28. Jiménez-Bremont, J. F., Marina, M., Guerrero-González, M. D., Rossi, F. R., Sánchez-Rangel, D., Rodríguez-Kessler, M., Ruiz, O. & Gárriz, A. 2014. Physiological and molecular implications of plant polyamine metabolism during biotic interactions. Frontiers in Plant Science, 5.

29. Joshi, K., Ahmed, S., Ge, L., Avestakh, A., Oloyede, B., Phuntumart, V., Kalinoski, A. & Morris, P. F. 2024. Spatial organization of putrescine synthesis in plants. Plant Sci, 349, 112232.

30. Joshi, V. & Fernie, A. R. 2017. Citrulline metabolism in plants. Amino Acids, 49, 1543–1559.

31. Kaur-Sawhney, R., Shih, L. M., Flores, H. E. & Galston, A. W. 1982. Relation of polyamine synthesis and titer to aging and senescence in oat leaves. Plant Physiol, 69, 405–10.

32. Kim, N. H., Kim, B. S. & Hwang, B. K. 2013. Pepper Arginine Decarboxylase Is Required for Polyamine and γ-Aminobutyric Acid Signaling in Cell Death and Defense Response. Plant Physiology, 162, 2067–2083.

33. Kwak, S. H. & Lee, S. H. 2001. The regulation of ornithine decarboxylase gene expression by sucrose and small upstream open reading frame in tomato (Mill). Plant and Cell Physiology, 42, 314–323.

34. Liang, J., Han, Q., Tan, Y., Ding, H. & Li, J. 2019. Current Advances on Structure-Function Relationships of Pyridoxal 5’-Phosphate-Dependent Enzymes. Front Mol Biosci, 6, 4.

35. Liu, J. H., Wang, W., Wu, H., Gong, X. Q. & Moriguchi, T. 2015. Polyamines function in stress tolerance: from synthesis to regulation. Frontiers in Plant Science, 6.

36. Lopez-Agudelo, J. C., Goh, F. J., Tchabashvili, S., Huang, Y. S., Huang, C. Y., Lee, K. T., Wang, Y. C., Wu, Y., Chang, H. X., Kuo, C. H., Lai, E. M. & Wu, C. H. 2025. Rhizobium rhizogenes A4-derived strains mediate hyper-eVicient transient gene expression in Nicotiana benthamiana and other solanaceous plants. Plant Biotechnol J.

37. Majumdar, R., Barchi, B., Turlapati, S. A., Gagne, M., Minocha, R., Long, S. & Minocha, S. C. 2016. Glutamate, Ornithine, Arginine, Proline, and Polyamine Metabolic Interactions: The Pathway Is Regulated at the Post-Transcriptional Level. Frontiers in Plant Science, 7.

38. Michael, A. J. 2016. Biosynthesis of polyamines and polyamine-containing molecules. Biochem J, 473, 2315–29.

39. Moormann, J., Heinemann, B. & Hildebrandt, T. M. 2022. News about amino acid metabolism in plant-microbe interactions. Trends Biochem Sci, 47, 839–850.

40. Napieraj, N., Janicka, M. & Reda, M. 2023. Interactions of Polyamines and Phytohormones in Plant Response to Abiotic Stress. Plants (Basel), 12.

41. Pal, M., Szalai, G., Gondor, O. K. & Janda, T. 2021. Unfinished story of polyamines: Role of conjugation, transport and light-related regulation in the polyamine metabolism in plants. Plant Sci, 308, 110923.

42. Paschalidis, K., Tsaniklidis, G., Wang, B. Q., Delis, C., Trantas, E., Loulakakis, K., Makky, M., Sarris, P. F., Ververidis, F. & Liu, J. H. 2019. The Interplay among Polyamines and Nitrogen in Plant Stress Responses. Plants-Basel, 8.

43. Poulin, R., Lu, L., Ackermann, B., Bey, P. & Pegg, A. E. 1992. Mechanism of the Irreversible Inactivation of Mouse Ornithine Decarboxylase by Alpha-Difluoromethylornithine - Characterization of Sequences at the Inhibitor and Coenzyme Binding-Sites. Journal of Biological Chemistry, 267, 150–158.

44. Ranawaka, B., An, J. Y., Lorenc, M. T., Jung, H. Y. T., Sulli, M., Aprea, G., Roden, S., Llaca, V., Hayashi, S., Asadyar, L., Leblanc, Z., Ahmed, Z., Naim, F., de Campos, S. B., Cooper, T., de Felippes, F. F., Dong, P. F., Zhong, S. L., Garcia-Carpintero, V., Orzaez, D., Dudley, K. J., Bombarely, A., Bally, J., Winefield, C., Giuliano, G. & Waterhouse, P. M. 2023. A multi-omic resource for fundamental research and biotechnology. Nature Plants. resource for fundamental research and biotechnology. Nature Plants.

45. Richards, F. J. & Coleman, R. G. 1952. Occurrence of putrescine in potassium-deficient barley. Nature, 170, 460.

46. Rossi, F. R., Romero, F. M., Ruiz, O. A., Marina, M. & Garriz, A. 2018. Phenotypic and Genotypic Characterization of Mutant Plants in Polyamine Metabolism Genes During Pathogenic Interactions. Methods Mol Biol, 1694, 405–416.

47. Sanchez-Lopez, J., Camanes, G., Flors, V., Vicent, C., Pastor, V., Vicedo, B., Cerezo, M. & Garcia-Agustin, P. 2009. Underivatized polyamine analysis in plant samples by ion pair LC coupled with electrospray tandem mass spectrometry. Plant Physiol Biochem, 47, 592–8.

48. Sandmeier, E., Hale, T. I. & Christen, P. 1994. Multiple evolutionary origin of pyridoxal-5’-phosphate-dependent amino acid decarboxylases. Eur J Biochem, 221, 997–1002.

49. Sivakumar, H. P., Sundararajan, S., Rajendran, V. & Ramalingam, S. 2022. Genome wide survey, and expression analysis of Ornithine decarboxylase gene associated with alkaloid biosynthesis in plants. Genomics, 114, 84–94.

50. Smith, T. A. 1970. Biosynthesis and Metabolism of Putrescine in Higher Plants. Annals of the New York Academy of Sciences, 171, 988-+.

51. Smith, T. A. & Richards, F. J. 1962. The biosynthesis of putrescine in higher plants and its relation to potassium nutrition. Biochem J, 84, 292–4.

52. Stephan, A., Hahn-Löbmann, S., Rosche, F., Buchholz, M., Giritch, A. & Gleba, Y. 2018. Simple Purification of -Produced Recombinant Colicins: High-Yield Recovery of Purified Proteins with Minimum Alkaloid Content Supports the Suitability of the Host for Manufacturing Food Additives. International Journal of Molecular Sciences, 19.

53. Upadhyay, R. K., Fatima, T., Handa, A. K. & Mattoo, A. K. 2020. Polyamines and Their Biosynthesis/Catabolism Genes Are DiVerentially Modulated in Response to Heat Versus Cold Stress in Tomato Leaves (Solanum lycopersicumL.). Cells, 9.

54. Urano, K., Hobo, T. & Shinozaki, K. 2005. Arabidopsis ADC genes involved in polyamine biosynthesis are essential for seed development. FEBS Lett, 579, 1557–64.

55. Winter, G., Todd, C. D., Trovato, M., Forlani, G. & Funck, D. 2015. Physiological implications of arginine metabolism in plants. Frontiers in Plant Science, 6.

56. Wittmann, J., Brancato, C., Berendzen, K. W. & Dreiseikelmann, B. 2016. Development of a tomato plant resistant to *Clavibacter michiganensis* using the endolysin gene of bacteriophage CMP1 as a transgene. Plant Pathology, 65, 496–502.

57. Wu, D., von roepenack-Lahaye, E., Buntru, M., de Lange, O., Schandry, N., Perez-Quintero, A. L., Weinberg, Z., Lowe-Power, T. M., Szurek, B., Michael, A. J., Allen, C., Schillberg, S. & Lahaye, T. 2019. A Plant Pathogen Type III EVector Protein Subverts Translational Regulation to Boost Host Polyamine Levels. Cell Host Microbe, 26, 638–649 e5.

58. Zierer, W., Hajirezaei, M. R., Eggert, K., Sauer, N., vonWirén, N. & Pommerrenig, B. 2016. Phloem-Specific Methionine Recycling Fuels Polyamine Biosynthesis in a Sulfur-Dependent Manner and Promotes Flower and Seed Development. Plant Physiology, 170, 790–806.

